# Leaf Mold Agar Facilitates Recovery of Soil Bacterial Diversity Beyond *Bacillus* Dominance

**DOI:** 10.1101/2025.05.13.653698

**Authors:** Fumiaki Tabuchi, Kazuhiro Mikami, Masaki Ishii, Jyunichiro Yasukawa, Masanobu Miyauchi, D.P.N. de Silva, Atsushi Miyashita

**Author notes:** Corresponding address: Atsushi Miyashita: Senior Assistant Professor; 359 Otsuka, Hachioji, Tokyo 192-0395, JAPAN.

## Abstract

Dominance of *Bacillus* species on conventional agar media often hampers the isolation of taxonomically diverse soil bacteria (i.e., “*Bacillus* dominance”). To overcome this *Bacillus* dominance, we developed a novel agar medium containing leaf mold extract (LME) and assessed its utility for isolating microorganisms from soil samples. For comparison, conventional yeast malt extract agar (YME agar) was used as a control. 16S rRNA gene sequencing revealed that colonies cultured on YME agar predominantly consisted of *Bacillus*-related bacteria, whereas LME agar enabled the growth of a wider variety of taxa, particularly actinomycetes such as *Streptomyces*. Notably, LME agar supported a higher diversity of soil bacterial isolates even from samples with a low abundance of *Bacillus*-related species. Among the 138 isolates recovered from LME agar, some either exhibited very low 16S rRNA gene sequence identity to known species in public databases, or failed to yield 16S rRNA amplicons with conventional universal primers, suggesting the presence of previously uncharacterized or novel taxa. Furthermore, one unidentified isolate failed to grow on standard nutrient media (YME, BHI, LB10, or TSB), but proliferated in liquid leaf mold extract, indicating a specific nutrient dependency. These findings demonstrate that LME agar can facilitate the isolation of a broader spectrum of soil microorganisms (including rare and previously uncultured species) not readily recoverable on common basal media. Incorporation of ecosystem-derived substrates into culture media may greatly enhance the discovery and study of novel microbial diversity from environmental samples.

**Importance:** Soil contains a vast number of microorganisms, many of which remain undiscovered due to limitations in standard laboratory methods. A major issue is the overgrowth of *Bacillus* species on conventional media, which suppresses the growth of other bacteria. To address this, we developed a new agar medium using leaf mold extract (LME) and found it successfully supported the growth of a much wider variety of soil bacteria, including rare species. Some of the isolated microbes could not be identified using standard genetic tools, and one unique strain only grew in LME-based media, indicating a special nutrient requirement. These findings show that using natural environmental components like leaf mold can help scientists grow and study bacteria that were previously missed. This approach can expand our understanding of microbial diversity and may lead to the discovery of new microbes with potential applications in agriculture, medicine, or environmental science.

## Introduction

Since the discovery of antibiotics, numerous pharmaceutically important compounds, including streptomycin, micafungin, ivermectin, tacrolimus, and doxorubicin, have been isolated from the culture supernatants of soil microorganisms (1–6). In recent years, novel antibiotics such as lysocin E, effective against Methicillin-resistant *Staphylococcus aureus* (MRSA), have also been discovered from soil-derived bacteria (7, 8). It is believed that among the microorganisms that have not yet been cultured, many may produce unknown compounds that could potentially serve as pharmaceutical candidates. However, the vast majority of environmental microbes remain uncultured and unexploited, representing an immense unreached source of potentially valuable bioactive compounds. Thus, developing innovative cultivation strategies or novel culture media is essential for accessing this hidden microbial diversity.

Soil represents one of the most microbially rich ecosystems, with an estimated 10^10^ to 10^11^ bacterial cells per gram (9). Environmental DNA analyses further reveal that soil harbors immense microbial diversity, much of which cannot currently be cultured by standard laboratory methods (10, 11). This “great plate count anomaly” has long hindered comprehensive study and bioprospecting of soil microbial communities (12).

Traditionally, microbes have been isolated from environmental samples by plating on nutrient-rich agar media (13). However, this approach often results in “*Bacillus* dominance”, where fast-growing bacteria (such as *Bacillus* spp.) dominate, suppressing the growth of slower-growing or more fastidious species and leading to reduced recoverable diversity. Several strategies have been explored to address this limitation, including the use of specialized agars (e.g., Soil Compost Agar at elevated temperatures to recover actinobacteria (14)), dilution of nutrient concentrations (15) (since rich nutrients can be toxic to some microorganisms (16, 17)), and supplementation with microbial supernatants to stimulate the growth of otherwise unculturable taxa (18). Despite these advances, a large fraction of environmental microbes remains inaccessible to conventional cultivation (19, 20).

To address this challenge, alternative media that better mimic natural environments are needed. In this study, we developed a new agar medium containing leaf mold extract (LME) and evaluated its effectiveness in cultivating diverse soil microorganisms. By comparing bacterial isolates obtained from LME agar and conventional yeast malt extract agar (YME), we assessed whether LME agar improves the recovery of phylogenetically diverse and previously uncultured or rare soil bacteria, as determined by 16S rRNA gene analysis.

## Materials and Methods

### Soil Sample Collection

Soil samples for this study were collected from three locations: 1) Musashino Central Park, Tokyo, Japan; 2) Abira Town, Hokkaido, Japan; and 3) Mount Jinba, Kanagawa, Japan. Each soil sample was collected using sterile plastic tubes (50 mL or 15 mL Falcon® tubes [Corning Life Sciences, Cat. Nos. 352098 and 352095, USA], or 2 mL Eppendorf tubes [Eppendorf SE, Cat. No. 0030120094, Germany]). For clarity, these samples are referred to throughout the manuscript as soil sample 1, soil sample 2, and soil sample 3, respectively.

### YME agar plates

The YME agar medium used in this study was prepared as follows: 4g of Yeast extract (Gibco, Cat. No., 211929, USA), 10g of Malt extract (Gibco, Cat. No., 218630, USA), 4g of D-glucose (Fujifilm, Cat. No., 049-31165, Japan), and 15g of Agar (Nacalai Tesque, Inc., Cat. No., 01028-85, Japan) were added to 1L of reverse osmosis (RO) water. This mixture was then autoclaved at 121°C for 15 minutes (Tomy, Cat. No., LSX-300, Japan). After autoclaving, 20 mL aliquots of the medium were poured into sterile plastic Petri dishes (AS ONE Corporation, Cat. No. 1-7484-01, Japan) and allowed to solidify prior to use for microbial cultivation.

### Leaf mold extract (LME) agar preparation

Leaf mold (product name: leaf mold with Bark, from a local supplier, Viva Home Hachioji, Japan) was used to prepare the medium. Leaf mold (approximately 500 g) was mixed with 2 L of Milli-Q water in a beaker, autoclaved at 121°C for 20 minutes, and then left to stand at room temperature overnight. The liquid extracted by heating was centrifuged (Hitachi, Cat. No., himac CR21E, Japan) at 9,300 g for 5 minutes and the supernatant was used as the leaf mold extract. The leaf mold extract was diluted 10-fold with pure water, and 1.5% agar was added. After autoclaving at 121°C for 15 minutes, 20 mL of the mixture was dispensed into plastic petri dishes to create the leaf mold agar plates (Fig. 1).

**Figure 1.**
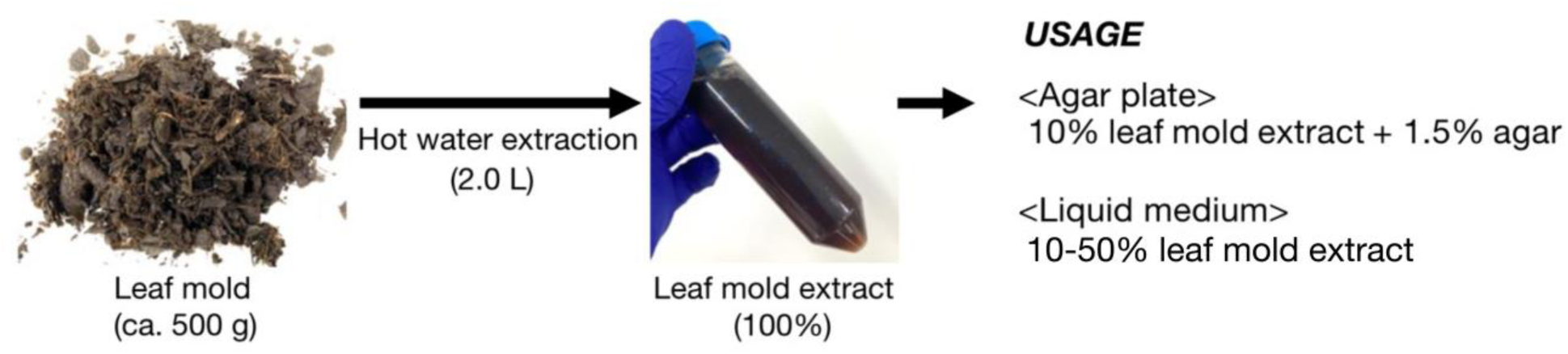
Preparation and application of leaf mold extract for microbial culture. Leaf mold (approximately 500 g) was subjected to hot water extraction with 2.0 L of Milli-Q water and autoclaved to obtain a concentrated (100%) leaf mold extract. The extract was used to prepare culture media as follows: for agar plates, it was diluted to 10% (v/v) and solidified with 1.5% agar; for liquid media, the extract was used at concentrations ranging from 10–50%.

### Bacterial culture and isolation

A spatula-full of each soil sample (the soil samples 1, 2, and 3) was suspended in 1 mL of sterile physiological saline and vortexed briefly. The suspension was serially diluted 10-fold and 100-fold. From each dilution, 100 μL aliquots were spread onto YME agar and LME agar plates, which were then incubated at 30°C for 3 days. In total, 116 colonies (sample 1: 68 strains; sample 2: 24 strains; sample 3: 24 strains) were isolated from YME agar, and 196 colonies (sample 1: 100 strains; sample 2: 48 strains; sample 3: 48 strains) were isolated from LME agar. Each isolate was subsequently restreaked onto fresh YME agar and LME agar plates and incubated at 30°C for several additional days to obtain purified single colonies.

### Amplification of DNA Sequences by Colony Direct PCR

Colony direct PCR was performed using single colonies isolated from each medium as templates. For amplification of 16S rRNA genes, two DNA polymerase systems were employed: Premix EX Taq™ Hot Start Version (TaKaRa Bio, Cat. No. RR006A, Japan) and KOD One® PCR Master Mix-Blue (TOYOBO, Cat. No. KMM-201, Japan). The primers used were 16S_8F (5’-AGAGTTTGATCCTGGCTCAG-3’) and 16S_1492R (5’-GGTTACCTTGTTACGACTT-3’). PCR with Premix EX Taq was performed with the following cycling conditions (modified from the manufacturer’s protocol): initial denaturation at 94°C for 10 min; 30 cycles of 98°C for 10 sec, annealing at an optimized temperature (T = 45, 50, 55, 60, or 65°C; determined for each strain) for 30 sec, and extension at 72°C for 90 sec. PCR with KOD One® PCR Master Mix-Blue followed the manufacturer’s standard protocol: 30 cycles of 98°C for 10 sec, 50°C for 5 sec, and 68°C for 10 sec. As a result, PCR products were successfully obtained from 100 isolates (sample 1: 59 strains; sample 2: 20 strains; sample 3: 21 strains) grown on YME agar and 161 isolates (sample 1: 81 strains; sample 2: 34 strains; sample 3: 46 strains) grown on LME agar.

### 16S rRNA Sequence Analysis

The PCR products obtained were treated and purified with ExoSAP IT^TM^ Express PCR Product Cleanup (Thermo Fisher Scientific, Cat. No., 75001.1.ML, USA) and NucleoSpin Gel and PCR Clean-up (Macherey-Nagel, Cat. No., 740609, Germany) according to the manual. Using the purified DNA as a template, DNA sequence analysis was conducted using the 16S_8F primer. In this study, sequence information of 16S rRNA was obtained from 93 strains (sample 1: 57 strains, sample 2: 18 strains, sample 3: 18 strains) grown on YME media and 138 strains (sample 1: 71 strains, sample 2: 28 strains, sample 3: 39 strains) grown on LME agar media. Isolates for which PCR amplification or DNA sequencing was unsuccessful were not included in the subsequent analyses.

The obtained sequences, approximately 800 bp in length, were subjected to genus-level identification by BLAST (Basic Local Alignment Search Tool) search. Due to the limited coverage of curated 16S rRNA gene databases for soil bacteria, sequence identification was performed using the standard NCBI (Natural Center for Biotechnology Information) database. Isolates with less than 98.7% sequence similarity to their closest match in the database were considered unidentified strains. Sequence lengths and detailed BLAST results for each isolate are summarized in Supplementary Tables S1 and S2.

### Molecular phylogenetic analysis

The phylogenetic tree analysis of 16S rRNA sequences was performed using the maximum likelihood method with the General Time Reversible model. The percentage of replicate trees in which the associated taxa clustered together is indicated at the branch nodes. For heuristic tree searching, initial trees were automatically obtained by applying the Neighbor-Join and BioNJ algorithms to a matrix of pairwise distances estimated by the maximum composite likelihood (MCL) approach. A gamma distribution was used to model evolutionary rate differences among sites, and the rate variation model allowed for some sites to be evolutionarily invariable. The phylogenetic tree is drawn to scale, with branch lengths measured as the number of substitutions per site. Evolutionary analyses were conducted using MEGA11. Sequence data from six isolates (1 isolate from sample 1, and 5 isolates from sample 3) were excluded from the analysis due to a lack of homologous and readable sequence regions.

### Preparation of Liquid Media and Bacterial Growth Assay

For the liquid culture of microorganisms, the following media were prepared and used.

*YME (Yeast Malt Extract):* This medium was composed of 4g Yeast extract, 10g Malt extract, and 4g D-glucose dissolved in 1L of reverse osmosis (RO) water. The solution was then autoclaved at 121°C for 15 minutes.

*BHI (Brain Heart Infusion):* This medium was prepared by dissolving 37g of Brain Heart Infusion powder (BD, Cat. No., 237500, USA) in 1L of RO water, followed by autoclaving at 121°C for 15 minutes.

*LB10 (Luria-Bertani medium):* This medium included 10g Tryptone (Gibco, Cat. No., 211705, USA), 5g Yeast extract, and 10g Sodium chloride (Fujifilm, Cat. No., 195-01663, Japan) dissolved in 1L of RO water. The solution was autoclaved at 121°C for 15 minutes.

*TSB (Tryptic Soy Broth):* This medium was prepared by dissolving 30g of Tryptic Soy Broth powder (BD, Cat. No., 211825, USA) in 1L of RO water and autoclaving at 121°C for 15 minutes.

*Leaf mold medium:* To evaluate whether bacterial growth could be consistently supported using different commercial leaf mold products, two additional leaf mold sources, ‘leaf mold’ (Tamiya Engei, Japan) and ‘homemade leaf mold’ (Akagi Engei, Japan), were used to prepare liquid media. For each product, 150 g of leaf mold was mixed with 500 mL of Milli-Q water in a beaker, autoclaved at 121°C for 15 minutes, and allowed to stand at room temperature overnight. The resulting extracts were diluted to final concentrations of 10% and 50% (v/v) and used as liquid media.

*Bacterial growth assay:* To assess the growth of bacterial isolates obtained from LME agar, colonies of a representative bacterial isolate (obtained using LME agar) were inoculated into each liquid medium and incubated at 30°C with shaking for 3 days. Comparable bacterial growth was observed in each medium, demonstrating that similar results could be obtained using leaf mold extracts from different commercial products.

## Results

### Distinct Colony Morphology on Different Media

After three days of incubation, clear differences were observed in colony morphology between YME agar and leaf mold extract agar (LME agar) plates inoculated with soil samples (Fig. 2). On YME agar, the majority of colonies were large, spreading, and frequently merged with adjacent colonies, typically corresponding to *Bacillus* species. In contrast, LME agar plates showed a greater number of small, well-separated colonies, many of which displayed morphologies consistent with actinomycetes, such as *Streptomyces*. These observations suggest that LME suppresses the overgrowth of fast-growing taxa and promotes the emergence of morphologically diverse colonies that are not readily recovered on conventional nutrient-rich media.

**Figure 2.**
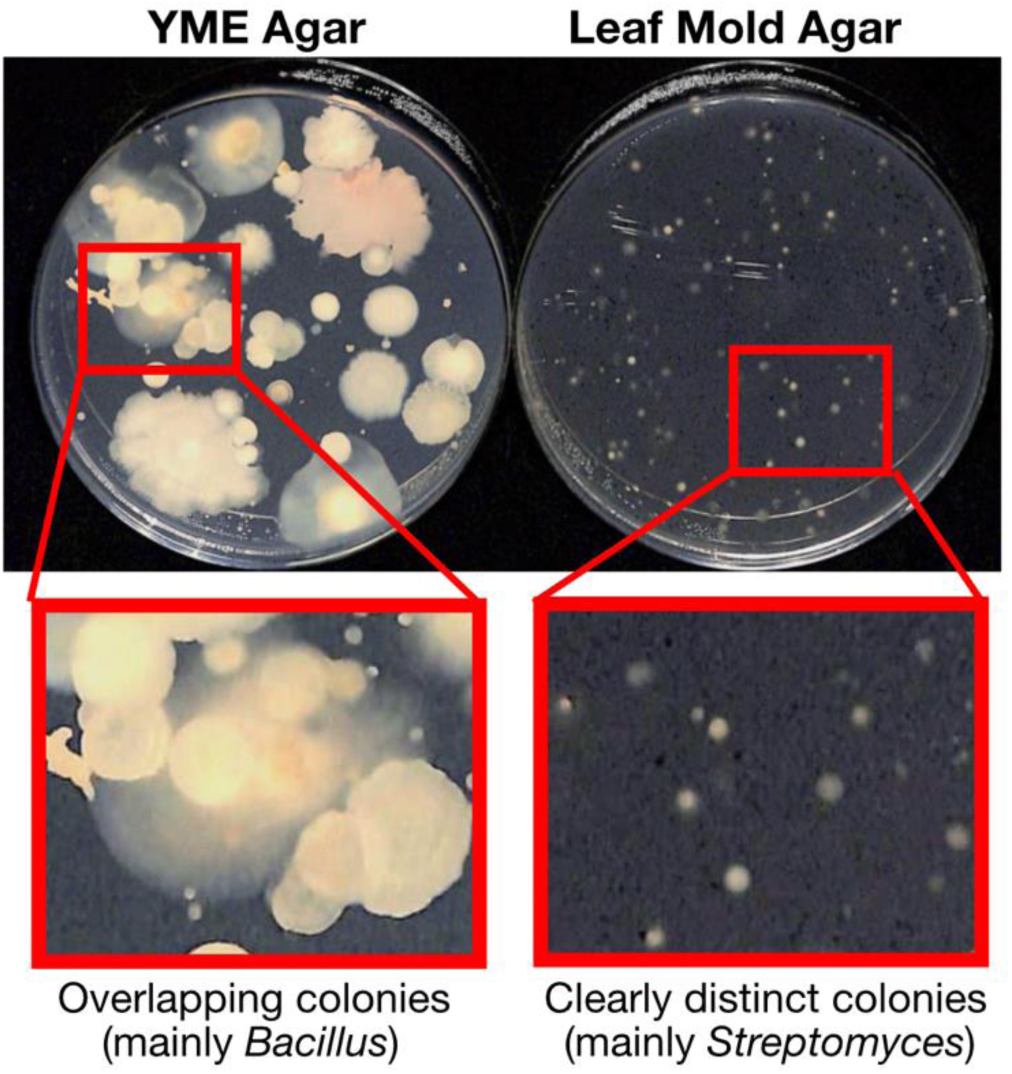
Colony morphology differences between YME agar and leaf mold agar. Representative images of colonies formed by soil microorganisms after incubation at 30°C for 3 days on conventional YME agar (left) and leaf mold agar (right). Magnified insets highlight differences: on YME agar, colonies are large and overlapping, primarily *Bacillus* species, while leaf mold agar promotes smaller, clearly separated colonies, predominantly *Streptomyces* species.

### Enhanced Phylogenetic Diversity on Leaf Mold Agar

To determine the taxonomic diversity of bacterial isolates, 16S rRNA gene sequencing was performed for 93 colonies isolated from YME agar and 138 from LME, derived from three independent soil samples. On YME agar, isolates were predominantly assigned to *Bacillus*-related genera (including *Bacillus*, *Priestia*, *Paenibacillus*, and *Neobacillus*), with a minor proportion (around 18%) belonging to the genus *Streptomyces* (Fig 3, Table S1). In contrast, LME agar supported a markedly more diverse community. For example, in soil sample 1, *Streptomyces* comprised 48% and *Rhizobium* 14% of the isolates on LME agar, along with representatives from *Actinacidiphila*, *Kitasatospora*, *Lysobacter*, and other genera, many of which were not detected on YME agar (Fig. 3, Table S2). A similar trend was observed with soil sample 2: LME agar yielded a dominance of *Streptomyces* (39%) and *Actinophytocola* (18%), whereas YME once again favored *Bacillus* and *Priestia*. Notably, in soil sample 3, where *Bacillus*-related species were scarce, LME agar still enabled the isolation of a broad range of actinomycetes and other taxa, highlighting the ability of LME agar to facilitate recovery of bacteria even from samples with inherently low *Bacillus* abundance. These results indicate that LME agar broadens the spectrum of culturable bacterial taxa relative to YME agar, with a particular enhancement for actinomycetes and other slow-growing or rare genera.

**Figure 3.**
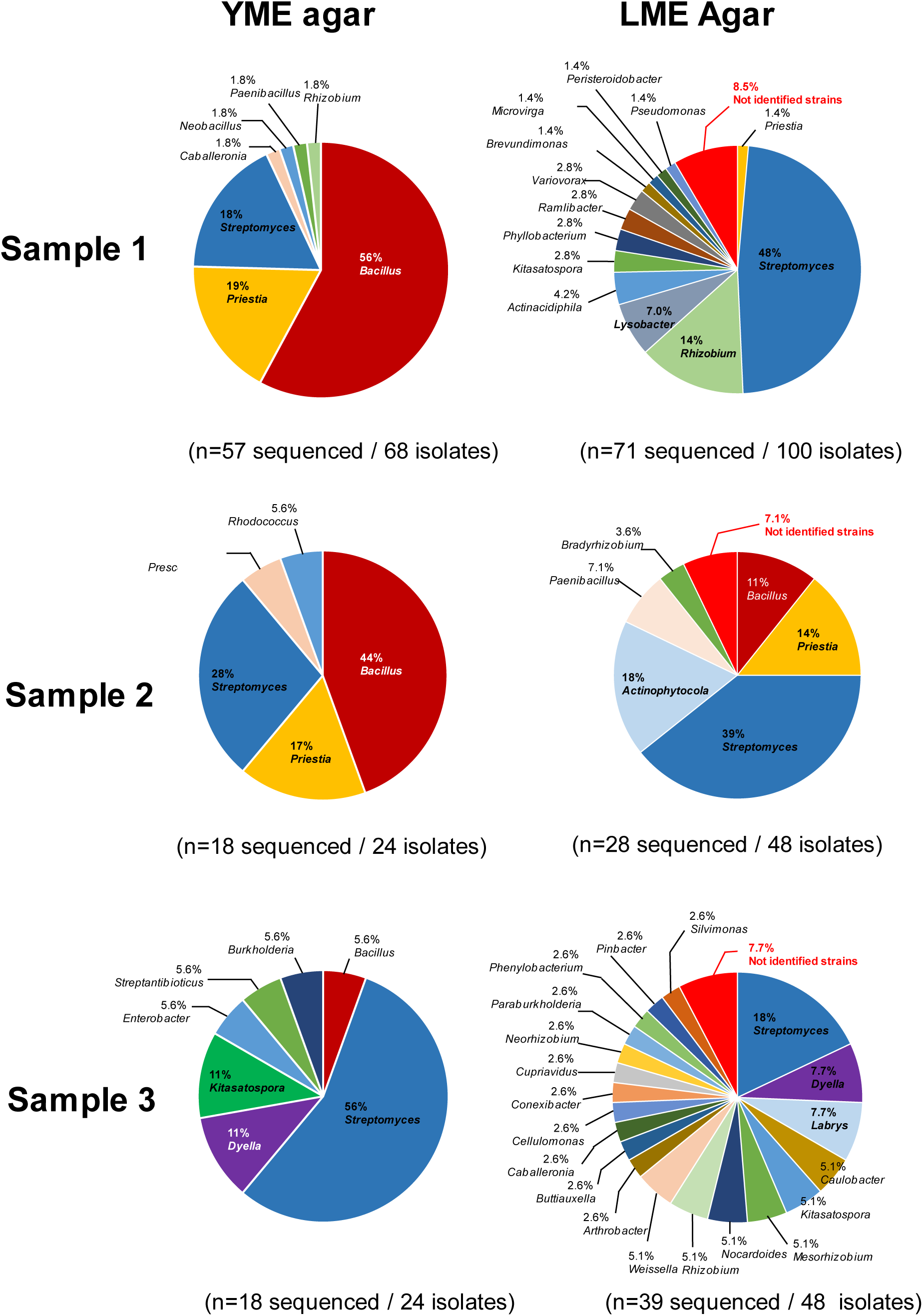
Comparison of microbial diversity cultivated on YME agar and leaf mold agar. Pie charts illustrate the distribution of bacterial genera isolated from three different soil samples on YME agar and leaf mold agar, as determined by 16S rRNA gene sequencing and NCBI database matching. For each soil sample, the genus-level diversity on YME (left) and leaf mold agar (right) is compared. Detailed isolate information is provided in Tables S1 and S2.

### Recovery of Potentially Novel and Unidentified Isolates

Among the colonies isolated from LME agar, several exhibited less than 98.7% 16S rRNA gene sequence identity to the closest species in the NCBI database (for example, a strain related to *Duganella* sp. at 98.69% identity), classifying them as unidentified or potentially novel species according to commonly accepted thresholds. In addition, some isolates from LME agar produced 16S rRNA amplicons that could not be reliably sequenced using standard universal primers, further suggesting taxonomic novelty. In contrast, nearly all isolates recovered from YME agar showed high sequence identity to described species. This finding demonstrates that incorporating ecosystem-derived substrates such as leaf mold can increase the probability of isolating uncharacterized or previously uncultured bacteria from environmental samples (Table S2).

### Phylogenetic Analysis Confirms Broader Lineage Access with LME agar

A maximum likelihood phylogenetic tree, constructed from partial 16S rRNA gene sequences of isolates from all three soil samples and both media, further demonstrated the contrasting spectrum of recovered bacteria (Fig. 4). YME agar-derived isolates clustered tightly within Bacillota (formerly Firmicutes), indicating low phylogenetic diversity. Conversely, LME agar-derived isolates were more widely distributed across the tree, encompassing a broad range of Actinomycetota and Proteobacteria, as well as several distinct clades that are separate from established groups. Notably, multiple LME agar isolates with low similarity to known taxa (indicated by a bold magenta label with a black “U” in the upper left corner) formed unique branches, supporting LME agar’s capability to isolate phylogenetically more distant microorganisms compared to YME medium. This phylogenetic assessment corroborates the genus-level findings and highlights LME agar as a tool to overcome the limitations of the “great plate count anomaly”.

**Figure 4.**
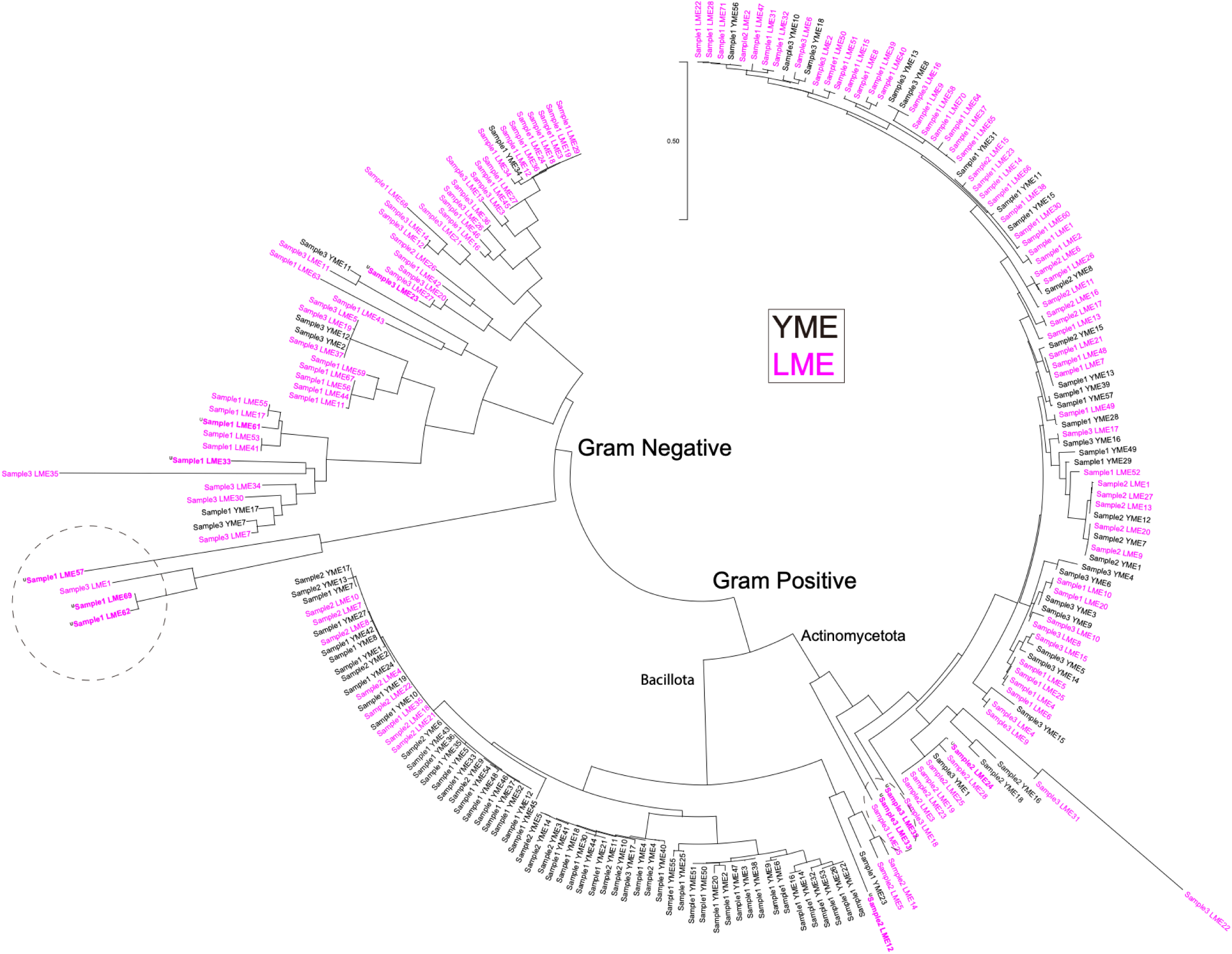
Phylogenetic tree of soil bacterial isolates cultured on YME agar and leaf mold agar. Maximum likelihood phylogenetic tree based on partial 16S rRNA gene sequences of bacteria isolated from three soil samples cultivated on YME agar (black) and leaf mold agar (magenta). Branches are annotated according to the isolation medium: “YME” (black) or “LME” (magenta). Isolates on LME agar with low similarity to known taxa were indicated by a bold magenta label with a black “U” in the upper left corner. Major taxonomic groups (including Bacillota and Actinomycetota) are indicated. The tree illustrates how isolates from leaf mold (LME) agar exhibit greater phylogenetic diversity, particularly within Actinomycetota, while Bacillota isolates are predominantly obtained from YME agar. A dashed circle highlights a distinct cluster of LME agar isolates, suggesting a potentially novel lineage. “Gram Negative” and “Gram Positive” clades are also annotated. Scale bar represents nucleotide substitutions per site.

### Specific Nutritional Requirements Demonstrated via Liquid Culture

To further probe the ecological breadth of microbes cultivable with LME agar, we investigated the liquid growth requirements of a strain isolated only on LME agar plates and classified as an unidentified bacterium. This strain failed to grow in four standard laboratory media (YME, BHI, LB10, TSB) but proliferated in liquid media prepared from leaf mold extract at both 10% and 50% concentrations, regardless of the leaf mold source (Fig. 5). This result underscores that certain soil bacteria may have specific dependencies on complex nutrient systems derived from natural environments, requirements that are not met by conventional laboratory media. It also highlights the importance of using ecosystem-mimetic substrates such as LME agar to access the full functional diversity of soil microbiota.

**Figure 5.**
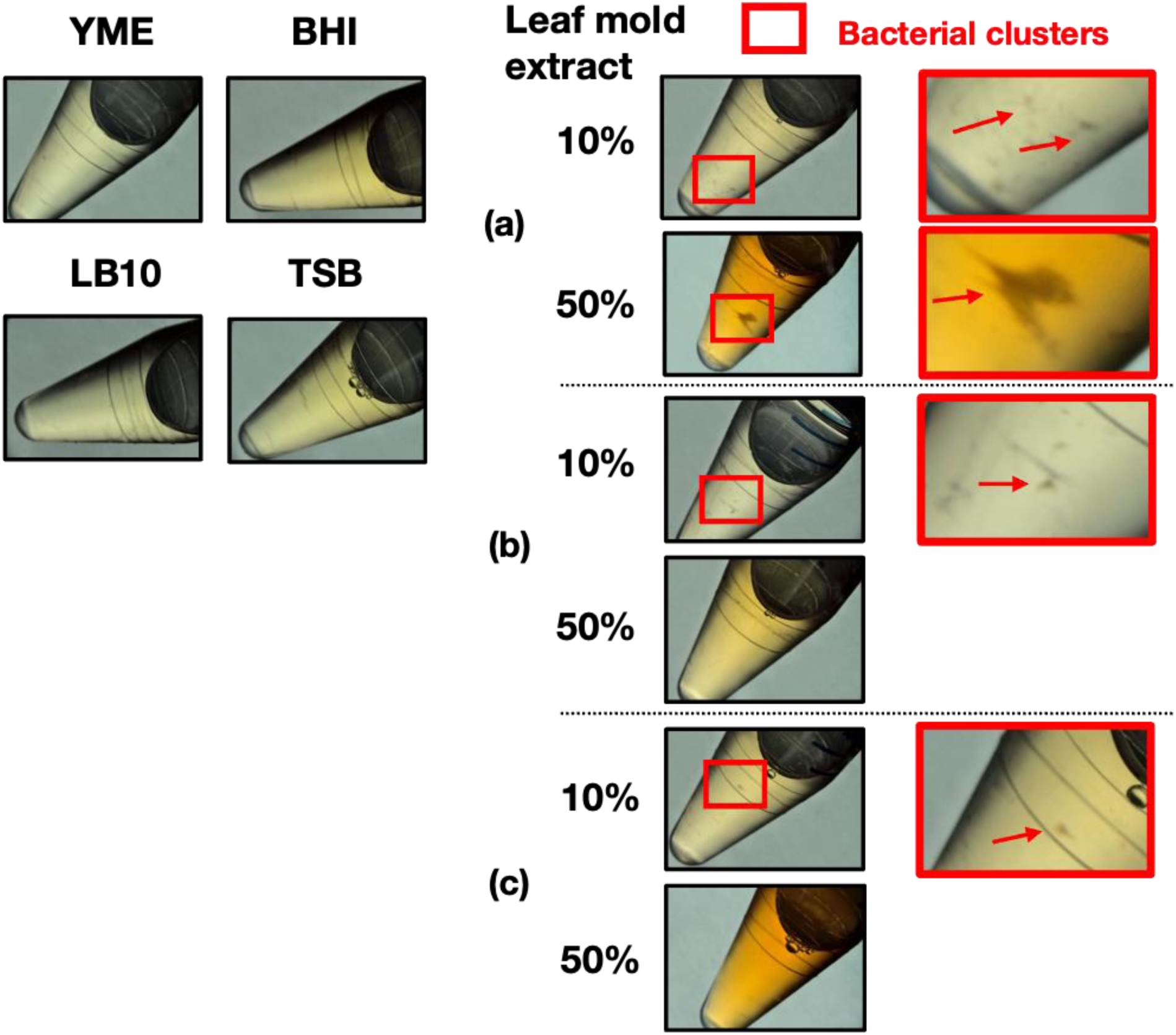
Growth of an unidentified bacterial isolate in nutrient broths and various leaf mold extract media. Colonies of an unidentified bacterial strain (most closely related to *Duganella* sp.) were inoculated in four standard nutrient broths (YME, BHI, LB10, and TSB) as well as in three different leaf mold extracts (a–c) at 10% and 50% concentrations. Tubes were incubated at 30°C for 3 days. No visible growth was observed in the nutrient broths, while bacterial clusters (highlighted by red squares and arrows) developed in all tested leaf mold extract media, as observed under a stereomicroscope. These results demonstrate the selective culturability of this isolate in leaf mold-based media.

## Discussion

In this study, we isolated bacteria from soil samples using agar media supplemented with leaf mold extract (LME agar) and analyzed their 16S rRNA gene sequences. We found that LME agar enabled the growth of bacterial colonies distinct from those observed on the basal YME medium by overcoming *Bacillus* dominance (Fig. 2, 3, and 4; Table S1 and S2). Consequently, LME agar elicited greater bacterial diversity, including the isolation of actinomycetes and rare bacterial taxa underrepresented on conventional media. Furthermore, some bacteria that were unable to grow on standard nutrient media successfully grew on liquid media supplemented with leaf mold extract (Fig. 5), suggesting the presence of growth-promoting components in the extract that facilitate the cultivation of soil bacteria that are otherwise underrepresented or overlooked on standard nutrient media.

The microbial community structures obtained from YME and LME agar were markedly different. YME agar predominantly yielded *Bacillus* species. In contrast, *Bacillus* growth was suppressed on LME agar, and actinomycetes such as *Streptomyces*, as well as other rare genera, were more frequently isolated. Phylogenetic analysis revealed that LME isolates primarily belonged to the Actinomycetota clade and exhibited greater phylogenetic diversity than those from YME (Fig. 4). In addition, several isolates from LME showed less than 98.7% sequence similarity to the closest matches in public databases, including some for which DNA sequencing of the 16S rRNA gene was unsuccessful, indicating the presence of potentially unidentified species (Table S1 and S2). These findings demonstrate that LME agar not only reduces *Bacillus* dominance but also facilitates access to a broader range of microbial taxa, including unidentified species that are often missed by conventional methods.

Several genera isolated on LME agar, including *Streptomyces* and *Lysobacter*, are known producers of valuable secondary metabolites such as streptomycin, cephamycin C, and lysocin E (2, 7, 8, 21). This highlights the practical significance of LME agar as a tool for microbial bioprospecting. The elevated diversity and frequency of such bioactive compound-producing bacteria on LME agar suggest its utility in drug discovery research. In our laboratory, we are developing screening systems using invertebrates such as silkworms (22–32). In addition to these screening systems, expanding microbial libraries with high diversity is essential for exploratory drug discovery. The use of LME agar to isolate a wide variety of soil bacteria supports the creation of such high-diversity microbial resources with potential for pharmaceutical applications.

Beyond solid media, certain strains that failed to grow on agar-based nutrient media were successfully cultivated in liquid culture containing leaf mold extract (Fig. 5). This supports the hypothesis that humic substances or other unknown components in the extract act as key growth factors. Although LME concentrations of 10% and 50% were tested in this study, further work is required to optimize cultivation parameters such as extract concentration, temperature, incubation time, and shaking speed. On the other hand, it is possible that LME lacked growth-inhibitory factors that are present in conventional nutrient media. Identifying and characterizing the growth-promoting or growth-inhibitory compounds may also clarify the molecular mechanisms underlying the improved cultivation of diverse soil microbes.

In relation to the cultivation of soil microbes, various improvements have been proposed to enhance the cultivation of environmental microbes, including nutrient dilution (15), supplementation with microbial supernatants (18), and *in situ* cultivation methods that mimic natural environmental conditions (33, 34). Among these, *in situ* cultivation has attracted attention for its potential to recover uncultured microbes, but it often requires specialized devices and poses challenges in maintaining environmental control. Despite such advances in cultivation technologies, many hard-to-culture microorganisms remain uncultivated. In contrast, the LME method demonstrated in this study is simple, inexpensive, reproducible using commercially available materials, and does not require special equipment or settings. Moreover, it resembles the approach used in humic acid–vitamin (HV) agar, which was developed for the selective isolation of diverse actinomycetes while suppressing the growth of non-actinomycete bacteria (35). This similarity may contribute to LME agar’s effectiveness in supporting actinomycete growth. It is necessary to purify and identify the factors in LME that explain the cultivation of diverse microorganisms.

Several isolates from LME agar exhibited less than 98.7% similarity in 16S rRNA gene sequences to the closest entries in public databases (36), including some for which DNA sequencing of the 16S rRNA gene was unsuccessful, suggesting that LME agar may serve as a valuable source for the isolation of potentially unidentified microbial species. Our phylogenetic analysis showed that some of these potentially novel isolates formed distinct clusters, further supporting their taxonomic novelty (Fig. 4). While some sequences were too short for definitive identification (Table S1 and S2), the use of LME agar clearly enabled access to taxonomically distinct strains. These potentially novel isolates may harbor unique genetic and metabolic traits, and continued exploration of such strains could lead to the discovery of novel bioactive compounds. Furthermore, expanding the application of LME agar to other environments, such as wetlands or deciduous forests, may further increase the diversity of cultivable microbes.

Overall, this study demonstrates that LME agar effectively circumvents the *Bacillus*-dominant bias of conventional media and enables the isolation of a more diverse array of soil bacteria, including uncultured and potentially novel species. By providing access to broader microbial diversity, LME agar significantly contributes to both basic microbial ecology and applied fields such as drug discovery. In particular, it offers a strong foundation for the development of diverse microbial libraries used in silkworms-based screening platforms.

## Author contribution

Conceptualization: AM, MI

Formal analysis: AM, FT, PS, MI, JY

Funding acquisition: AM

Investigation: AM, KM, MM, FT, JY

Project administration: AM, MI

Resources: AM

Supervision: AM

Visualization: AM, FT, MM, MI

Writing – original draft: AM, FT

Writing – review & editing: AM, KM, MI, PS, JY, FT

## Acknowledgment

This work was supported by JSPS KAKENHI (Grant# 22K15461, A.M.), grants from the Project of the NARO Bio-oriented Technology Research Advancement Institution (Research program on development of innovative technology) (Grant# 05001a1 and a1a2, A.M.), and Teikyo University Team Resarch Grant (Grant# 22-24, A.M.).

## Supplemental Material

**Table S1.**
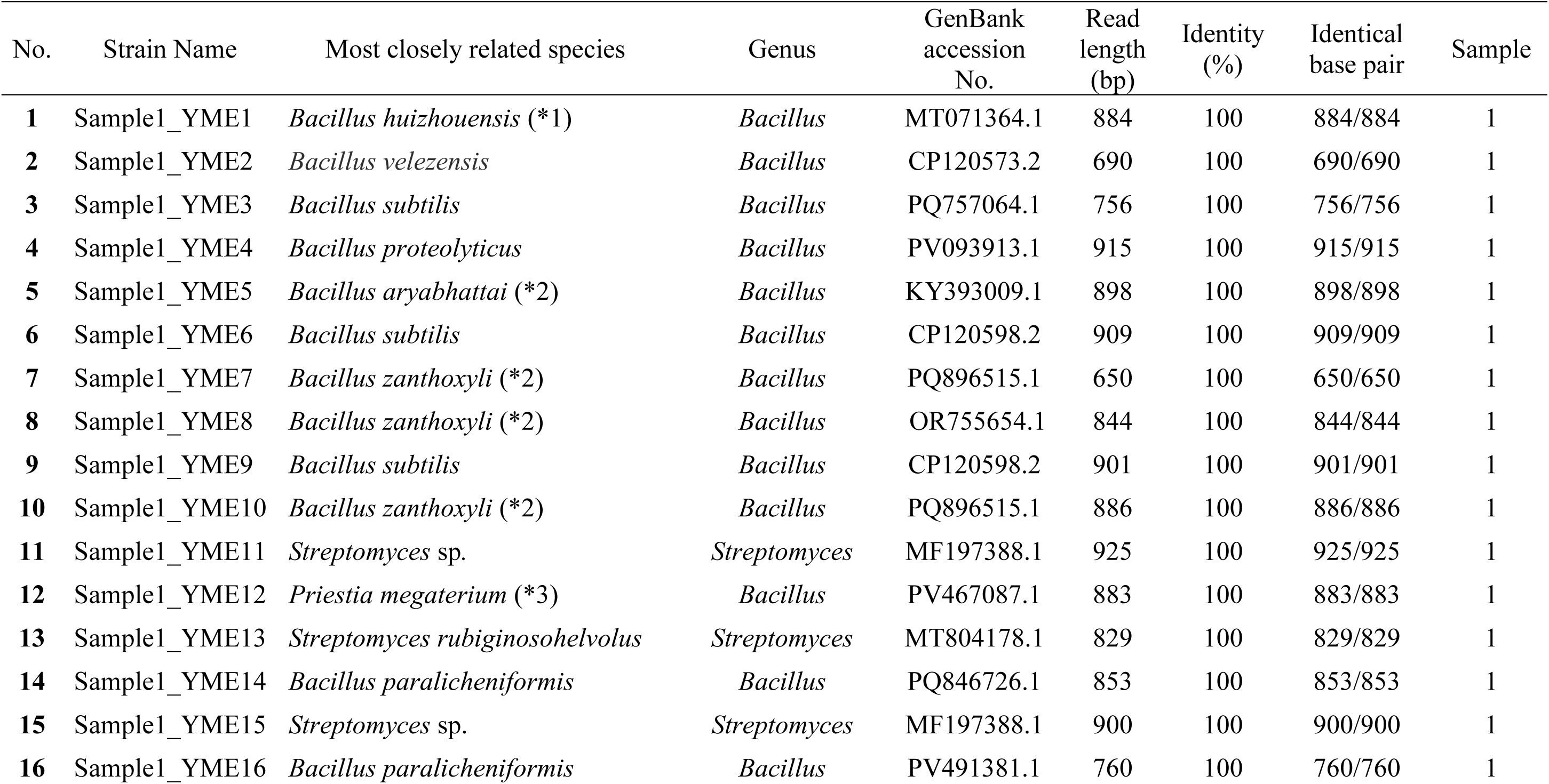
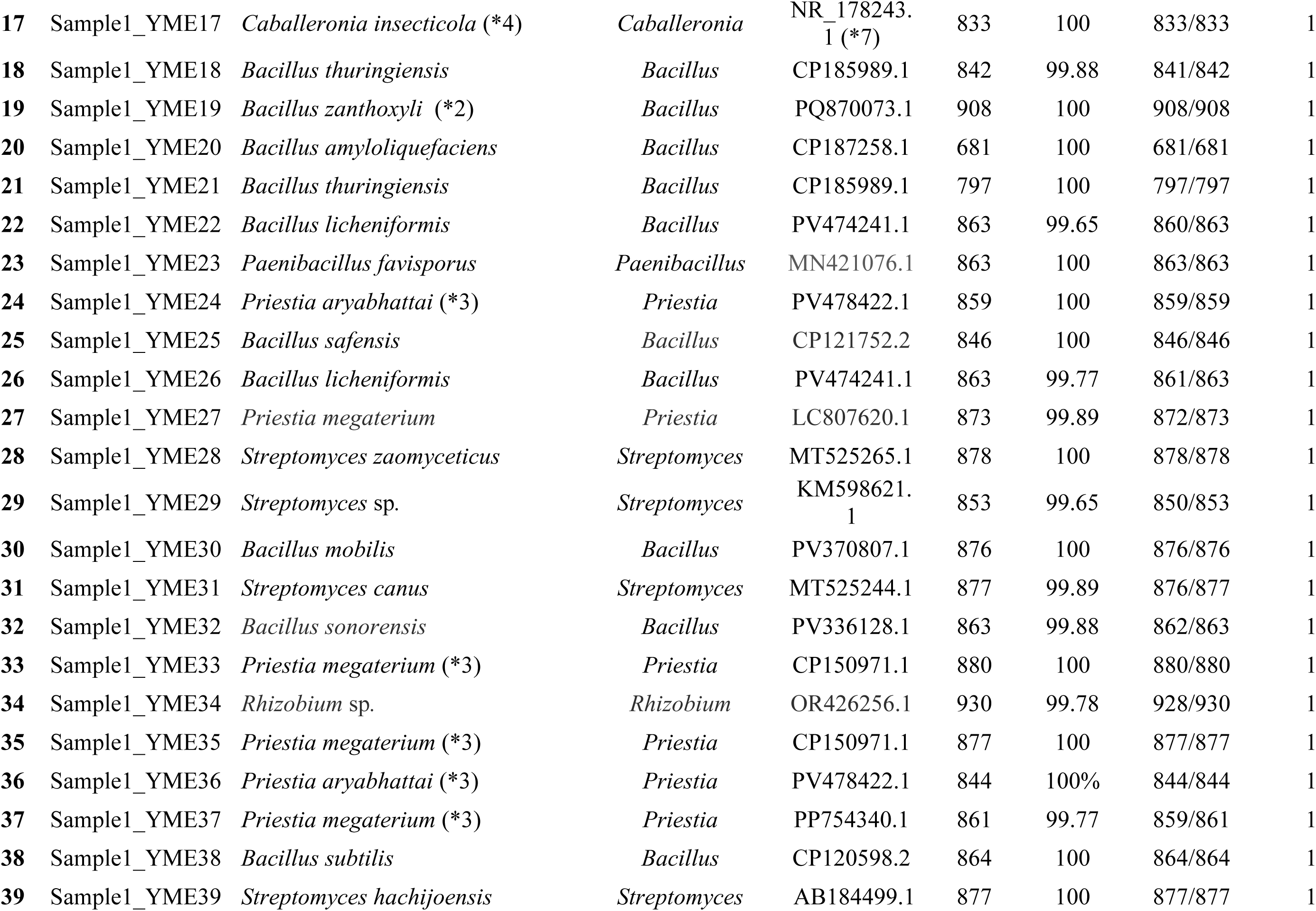
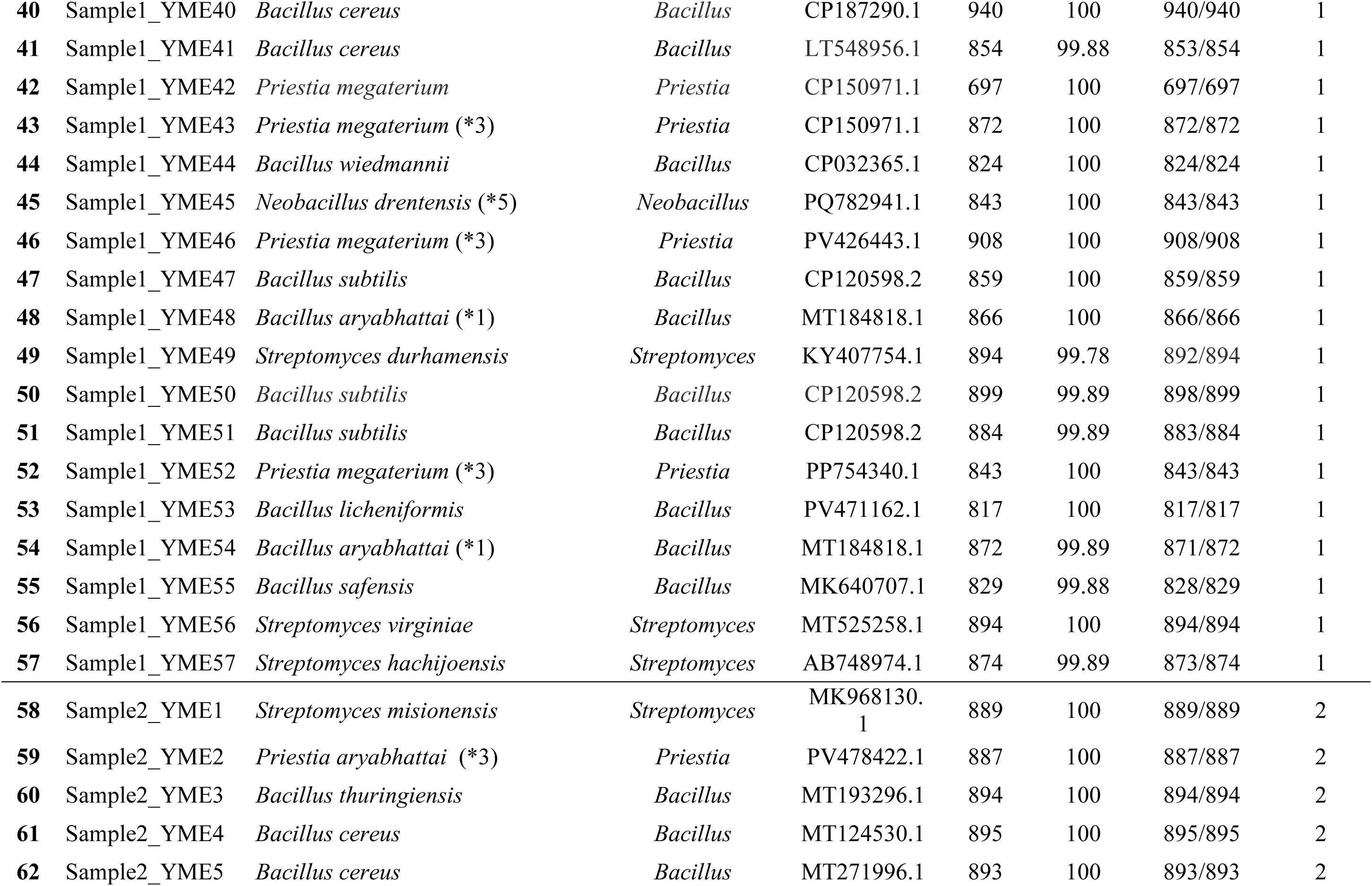
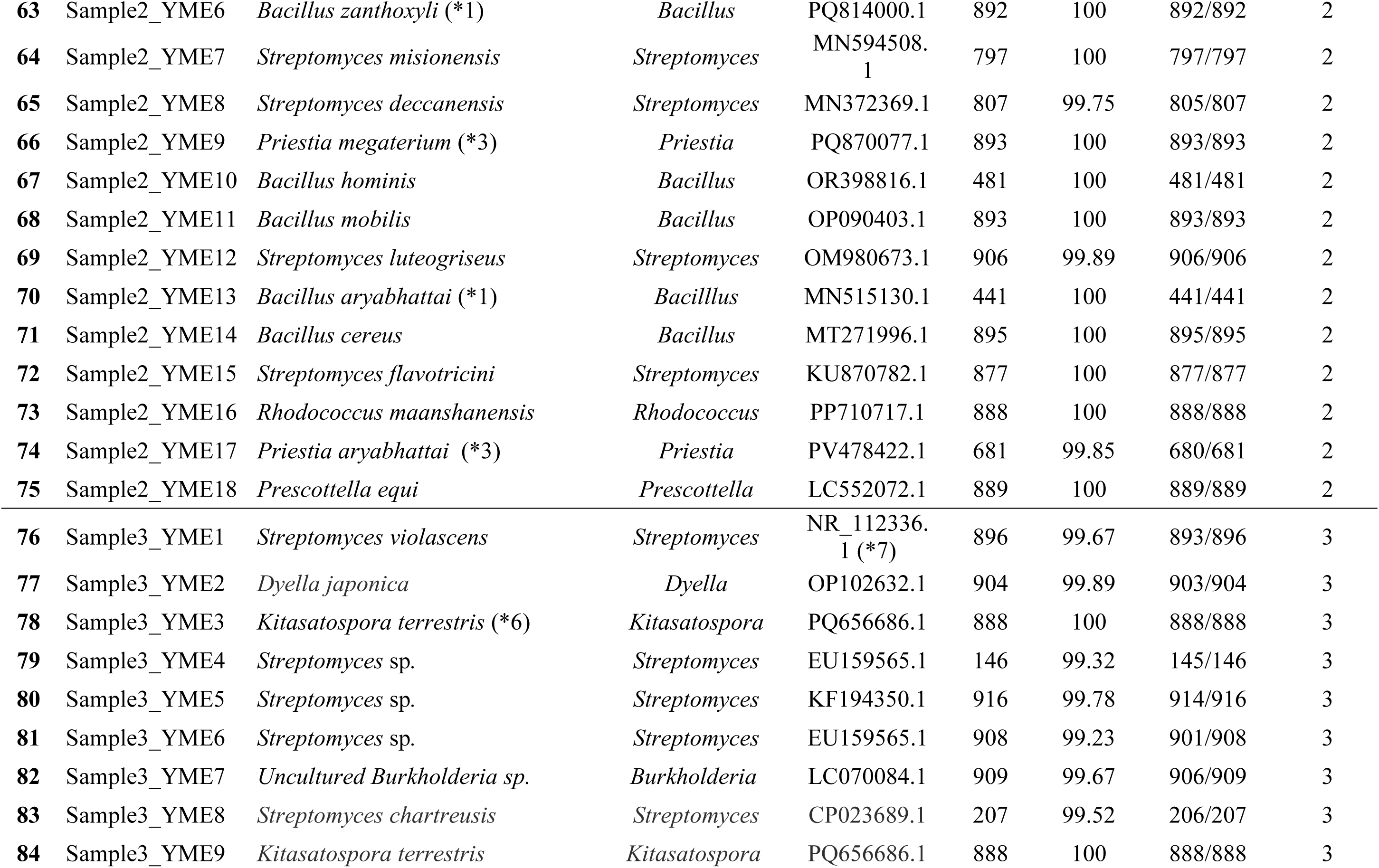
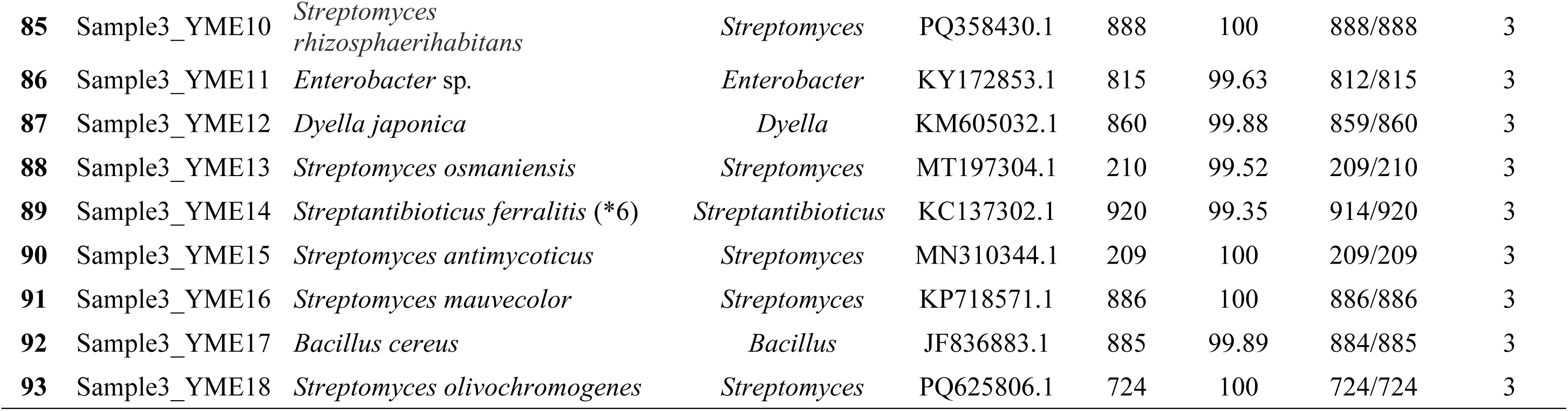
List of Bacteria Grown on YME Agar Medium The 16S rRNA sequences of 93 bacterial strains (sample 1: 57, sample 2: 18, sample 3: 18) grown on YME agar were analyzed and compared with the NCBI standard database. The closest related species in the database, their genus, GenBank accession number, read length, match rate, and number of matching basepairs are summarized in this table. Isolates marked with an asterisk (*) indicate that there were multiple species with the same match rate in the database for the 16S rRNA analysis conducted in this study. The respective species are as follows: *1: *Priestia megaterium*, *2: *Priestia aryabhattai*, *3: *Bacillus zanthoxyli*, *4: *Burkholderia* sp., *5: *Bacillus arbutinivorans*., *6: *Streptomyces* sp.. In cases where an NCBI Reference Sequence was shown instead of a GenBank accession number in the NCBI database, an asterisk followed by the number 7 (*7) was added. To draw the pie chart in Figure 3, the genus names listed in the table were used.

| No. | Strain Name | Most closely related species | Genus | GenBank<br>accession<br>No. | Read<br>length<br>(bp) | Identity<br>(%) | Identical<br>base pair | Sample |
| --- | --- | --- | --- | --- | --- | --- | --- | --- |
| 1 | Sample1_YME1 | *Bacillus huizhouensis* (*1) | *Bacillus* | MT071364.1 | 884 | 100 | 884/884 | 1 |
| 2 | Sample1_YME2 | *Bacillus velezensis* | *Bacillus* | CP120573.2 | 690 | 100 | 690/690 | 1 |
| 3 | Sample1_YME3 | *Bacillus subtilis* | *Bacillus* | PQ757064.1 | 756 | 100 | 756/756 | 1 |
| 4 | Sample1_YME4 | *Bacillus proteolyticus* | *Bacillus* | PV093913.1 | 915 | 100 | 915/915 | 1 |
| 5 | Sample1_YME5 | *Bacillus aryabhattai* (*2) | *Bacillus* | KY393009.1 | 898 | 100 | 898/898 | 1 |
| 6 | Sample1_YME6 | *Bacillus subtilis* | *Bacillus* | CP120598.2 | 909 | 100 | 909/909 | 1 |
| 7 | Sample1_YME7 | *Bacillus zanthoxyli* (*2) | *Bacillus* | PQ896515.1 | 650 | 100 | 650/650 | 1 |
| 8 | Sample1_YME8 | *Bacillus zanthoxyli* (*2) | *Bacillus* | OR755654.1 | 844 | 100 | 844/844 | 1 |
| 9 | Sample1_YME9 | *Bacillus subtilis* | *Bacillus* | CP120598.2 | 901 | 100 | 901/901 | 1 |
| 10 | Sample1_YME10 | *Bacillus zanthoxyli* (*2) | *Bacillus* | PQ896515.1 | 886 | 100 | 886/886 | 1 |
| 11 | Sample1_YME11 | *Streptomyces* sp. | *Streptomyces* | MF197388.1 | 925 | 100 | 925/925 | 1 |
| 12 | Sample1_YME12 | *Priestia megaterium* (*3) | *Bacillus* | PV467087.1 | 883 | 100 | 883/883 | 1 |
| 13 | Sample1_YME13 | *Streptomyces rubiginosohelvolus* | *Streptomyces* | MT804178.1 | 829 | 100 | 829/829 | 1 |
| 14 | Sample1_YME14 | *Bacillus paralicheniformis* | *Bacillus* | PQ846726.1 | 853 | 100 | 853/853 | 1 |
| 15 | Sample1_YME15 | *Streptomyces* sp. | *Streptomyces* | MF197388.1 | 900 | 100 | 900/900 | 1 |
| 16 | Sample1_YME16 | *Bacillus paralicheniformis* | *Bacillus* | PV491381.1 | 760 | 100 | 760/760 | 1 |
| 17 | Sample1_YME17 | *Caballeronia insecticola* (*4) | *Caballeronia* | NR_178243.1 (*7) | 833 | 100 | 833/833 | 1 |
| 18 | Sample1_YME18 | *Bacillus thuringiensis* | *Bacillus* | CP185989.1 | 842 | 99.88 | 841/842 | 1 |
| 19 | Sample1_YME19 | *Bacillus zanthoxyli* (*2) | *Bacillus* | PQ870073.1 | 908 | 100 | 908/908 | 1 |
| 20 | Sample1_YME20 | *Bacillus amyloliquefaciens* | *Bacillus* | CP187258.1 | 681 | 100 | 681/681 | 1 |
| 21 | Sample1_YME21 | *Bacillus thuringiensis* | *Bacillus* | CP185989.1 | 797 | 100 | 797/797 | 1 |
| 22 | Sample1_YME22 | *Bacillus licheniformis* | *Bacillus* | PV474241.1 | 863 | 99.65 | 860/863 | 1 |
| 23 | Sample1_YME23 | *Paenibacillus favisporus* | *Paenibacillus* | MN421076.1 | 863 | 100 | 863/863 | 1 |
| 24 | Sample1_YME24 | *Priestia aryabhattai* (*3) | *Priestia* | PV478422.1 | 859 | 100 | 859/859 | 1 |
| 25 | Sample1_YME25 | *Bacillus safensis* | *Bacillus* | CP121752.2 | 846 | 100 | 846/846 | 1 |
| 26 | Sample1_YME26 | *Bacillus licheniformis* | *Bacillus* | PV474241.1 | 863 | 99.77 | 861/863 | 1 |
| 27 | Sample1_YME27 | *Priestia megaterium* | *Priestia* | LC807620.1 | 873 | 99.89 | 872/873 | 1 |
| 28 | Sample1_YME28 | *Streptomyces zaomyceticus* | *Streptomyces* | MT525265.1 | 878 | 100 | 878/878 | 1 |
| 29 | Sample1_YME29 | *Streptomyces* sp. | *Streptomyces* | KM598621.1 | 853 | 99.65 | 850/853 | 1 |
| 30 | Sample1_YME30 | *Bacillus mobilis* | *Bacillus* | PV370807.1 | 876 | 100 | 876/876 | 1 |
| 31 | Sample1_YME31 | *Streptomyces canus* | *Streptomyces* | MT525244.1 | 877 | 99.89 | 876/877 | 1 |
| 32 | Sample1_YME32 | *Bacillus sonorensis* | *Bacillus* | PV336128.1 | 863 | 99.88 | 862/863 | 1 |
| 33 | Sample1_YME33 | *Priestia megaterium* (*3) | *Priestia* | CP150971.1 | 880 | 100 | 880/880 | 1 |
| 34 | Sample1_YME34 | *Rhizobium* sp. | *Rhizobium* | OR426256.1 | 930 | 99.78 | 928/930 | 1 |
| 35 | Sample1_YME35 | *Priestia megaterium* (*3) | *Priestia* | CP150971.1 | 877 | 100 | 877/877 | 1 |
| 36 | Sample1_YME36 | *Priestia aryabhattai* (*3) | *Priestia* | PV478422.1 | 844 | 100% | 844/844 | 1 |
| 37 | Sample1_YME37 | *Priestia megaterium* (*3) | *Priestia* | PP754340.1 | 861 | 99.77 | 859/861 | 1 |
| 38 | Sample1_YME38 | *Bacillus subtilis* | *Bacillus* | CP120598.2 | 864 | 100 | 864/864 | 1 |
| 39 | Sample1_YME39 | *Streptomyces hachijoensis* | *Streptomyces* | AB184499.1 | 877 | 100 | 877/877 | 1 |
| 40 | Sample1_YME40 | *Bacillus cereus* | *Bacillus* | CP187290.1 | 940 | 100 | 940/940 | 1 |
| 41 | Sample1_YME41 | *Bacillus cereus* | *Bacillus* | LT548956.1 | 854 | 99.88 | 853/854 | 1 |
| 42 | Sample1_YME42 | *Priestia megaterium* | *Priestia* | CP150971.1 | 697 | 100 | 697/697 | 1 |
| 43 | Sample1_YME43 | *Priestia megaterium* (*3) | *Priestia* | CP150971.1 | 872 | 100 | 872/872 | 1 |
| 44 | Sample1_YME44 | *Bacillus wiedmannii* | *Bacillus* | CP032365.1 | 824 | 100 | 824/824 | 1 |
| 45 | Sample1_YME45 | *Neobacillus drentensis* (*5) | *Neobacillus* | PQ782941.1 | 843 | 100 | 843/843 | 1 |
| 46 | Sample1_YME46 | *Priestia megaterium* (*3) | *Priestia* | PV426443.1 | 908 | 100 | 908/908 | 1 |
| 47 | Sample1_YME47 | *Bacillus subtilis* | *Bacillus* | CP120598.2 | 859 | 100 | 859/859 | 1 |
| 48 | Sample1_YME48 | *Bacillus aryabhattai* (*1) | *Bacillus* | MT184818.1 | 866 | 100 | 866/866 | 1 |
| 49 | Sample1_YME49 | *Streptomyces durhamensis* | *Streptomyces* | KY407754.1 | 894 | 99.78 | 892/894 | 1 |
| 50 | Sample1_YME50 | *Bacillus subtilis* | *Bacillus* | CP120598.2 | 899 | 99.89 | 898/899 | 1 |
| 51 | Sample1_YME51 | *Bacillus subtilis* | *Bacillus* | CP120598.2 | 884 | 99.89 | 883/884 | 1 |
| 52 | Sample1_YME52 | *Priestia megaterium* (*3) | *Priestia* | PP754340.1 | 843 | 100 | 843/843 | 1 |
| 53 | Sample1_YME53 | *Bacillus licheniformis* | *Bacillus* | PV471162.1 | 817 | 100 | 817/817 | 1 |
| 54 | Sample1_YME54 | *Bacillus aryabhattai* (*1) | *Bacillus* | MT184818.1 | 872 | 99.89 | 871/872 | 1 |
| 55 | Sample1_YME55 | *Bacillus safensis* | *Bacillus* | MK640707.1 | 829 | 99.88 | 828/829 | 1 |
| 56 | Sample1_YME56 | *Streptomyces virginiae* | *Streptomyces* | MT525258.1 | 894 | 100 | 894/894 | 1 |
| 57 | Sample1_YME57 | *Streptomyces hachijoensis* | *Streptomyces* | AB748974.1 | 874 | 99.89 | 873/874 | 1 |
| 58 | Sample2_YME1 | *Streptomyces misionensis* | *Streptomyces* | MK968130.1 | 889 | 100 | 889/889 | 2 |
| 59 | Sample2_YME2 | *Priestia aryabhattai* (*3) | *Priestia* | PV478422.1 | 887 | 100 | 887/887 | 2 |
| 60 | Sample2_YME3 | *Bacillus thuringiensis* | *Bacillus* | MT193296.1 | 894 | 100 | 894/894 | 2 |
| 61 | Sample2_YME4 | *Bacillus cereus* | *Bacillus* | MT124530.1 | 895 | 100 | 895/895 | 2 |
| 62 | Sample2_YME5 | *Bacillus cereus* | *Bacillus* | MT271996.1 | 893 | 100 | 893/893 | 2 |
| 63 | Sample2_YME6 | *Bacillus zanthoxyli* (*1) | *Bacillus* | PQ814000.1 | 892 | 100 | 892/892 | 2 |
| 64 | Sample2_YME7 | *Streptomyces misionensis* | *Streptomyces* | MN594508.1 | 797 | 100 | 797/797 | 2 |
| 65 | Sample2_YME8 | *Streptomyces deccanensis* | *Streptomyces* | MN372369.1 | 807 | 99.75 | 805/807 | 2 |
| 66 | Sample2_YME9 | *Priestia megaterium* (*3) | *Priestia* | PQ870077.1 | 893 | 100 | 893/893 | 2 |
| 67 | Sample2_YME10 | *Bacillus hominis* | *Bacillus* | OR398816.1 | 481 | 100 | 481/481 | 2 |
| 68 | Sample2_YME11 | *Bacillus mobilis* | *Bacillus* | OP090403.1 | 893 | 100 | 893/893 | 2 |
| 69 | Sample2_YME12 | *Streptomyces luteogriseus* | *Streptomyces* | OM980673.1 | 906 | 99.89 | 906/906 | 2 |
| 70 | Sample2_YME13 | *Bacillus aryabhattai* (*1) | *Bacillus* | MN515130.1 | 441 | 100 | 441/441 | 2 |
| 71 | Sample2_YME14 | *Bacillus cereus* | *Bacillus* | MT271996.1 | 895 | 100 | 895/895 | 2 |
| 72 | Sample2_YME15 | *Streptomyces flavotricini* | *Streptomyces* | KU870782.1 | 877 | 100 | 877/877 | 2 |
| 73 | Sample2_YME16 | *Rhodococcus maanshanensis* | *Rhodococcus* | PP710717.1 | 888 | 100 | 888/888 | 2 |
| 74 | Sample2_YME17 | *Priestia aryabhattai* (*3) | *Priestia* | PV478422.1 | 681 | 99.85 | 680/681 | 2 |
| 75 | Sample2_YME18 | *Prescottella equi* | *Prescottella* | LC552072.1 | 889 | 100 | 889/889 | 2 |
| 76 | Sample3_YME1 | *Streptomyces violascens* | *Streptomyces* | NR_112336.1 (*7) | 896 | 99.67 | 893/896 | 3 |
| 77 | Sample3_YME2 | *Dyella japonica* | *Dyella* | OP102632.1 | 904 | 99.89 | 903/904 | 3 |
| 78 | Sample3_YME3 | *Kitasatospora terrestris* (*6) | *Kitasatospora* | PQ656686.1 | 888 | 100 | 888/888 | 3 |
| 79 | Sample3_YME4 | *Streptomyces* sp. | *Streptomyces* | EU159565.1 | 146 | 99.32 | 145/146 | 3 |
| 80 | Sample3_YME5 | *Streptomyces* sp. | *Streptomyces* | KF194350.1 | 916 | 99.78 | 914/916 | 3 |
| 81 | Sample3_YME6 | *Streptomyces* sp. | *Streptomyces* | EU159565.1 | 908 | 99.23 | 901/908 | 3 |
| 82 | Sample3_YME7 | *Uncultured Burkholderia* sp. | *Burkholderia* | LC070084.1 | 909 | 99.67 | 906/909 | 3 |
| 83 | Sample3_YME8 | *Streptomyces chartreusis* | *Streptomyces* | CP023689.1 | 207 | 99.52 | 206/207 | 3 |
| 84 | Sample3_YME9 | *Kitasatospora terrestris* | *Kitasatospora* | PQ656686.1 | 888 | 100 | 888/888 | 3 |
| 85 | Sample3_YME10 | *Streptomyces rhizosphaerihabitans* | *Streptomyces* | PQ358430.1 | 888 | 100 | 888/888 | 3 |
| 86 | Sample3_YME11 | *Enterobacter* sp. | *Enterobacter* | KY172853.1 | 815 | 99.63 | 812/815 | 3 |
| 87 | Sample3_YME12 | *Dyella japonica* | *Dyella* | KM605032.1 | 860 | 99.88 | 859/860 | 3 |
| 88 | Sample3_YME13 | *Streptomyces osmaniensis* | *Streptomyces* | MT197304.1 | 210 | 99.52 | 209/210 | 3 |
| 89 | Sample3_YME14 | *Streptantibioticus ferralitis* (*6) | *Streptantibioticus* | KC137302.1 | 920 | 99.35 | 914/920 | 3 |
| 90 | Sample3_YME15 | *Streptomyces antimycoticus* | *Streptomyces* | MN310344.1 | 209 | 100 | 209/209 | 3 |
| 91 | Sample3_YME16 | *Streptomyces mauvecolor* | *Streptomyces* | KP718571.1 | 886 | 100 | 886/886 | 3 |
| 92 | Sample3_YME17 | *Bacillus cereus* | *Bacillus* | JF836883.1 | 885 | 99.89 | 884/885 | 3 |
| 93 | Sample3_YME18 | *Streptomyces olivochromogenes* | *Streptomyces* | PQ625806.1 | 724 | 100 | 724/724 | 3 |

**Table S2.** List of Bacteria Grown on Leaf Mold Agar Medium The 16S rRNA sequences of 138 bacterial strains (sample 1: 71, sample 2: 28, sample 3: 39) grown on leaf mold agar were analyzed and matched with the NCBI rRNA sequence database (A: 127 identified species, B: 11 unidentified species). The species identified as the closest relative in the database, their genus, GenBank accession number, read length, match rate, and number of matching base pairs are compiled in this table as Identified isolates. Strains with a match rate less than 98.7% to the closest related strain in the database were designated as unidentified species and compiled as Unidentified isolates. Isolates marked with an asterisk (*) indicate that there were multiple species with the same match rate in the database for the 16S rRNA analysis conducted in this study. The respective species are as follows: *1: *Streptomyces* sp., *2: *Actinacidphila glacinigra*, *3: *Bacillus zanthoxyli*, *4: *Priestia megaterium.* However, the genus names listed in the table were used when creating the pie chart in Figure 3. In cases where an NCBI Reference Sequence was shown instead of a GenBank accession number in the NCBI database, an asterisk followed by the number 5 (*5) was added. The bacterial species shown in the “Most closely related species” column of list B were indicated in the database as the closest matches to the respective unidentified isolates.

| No. | Strain Name | Most closely related species | Genus | GenBank accession No. | Read length (bp) | Identity (%) | Identical base pair | Sample |
| --- | --- | --- | --- | --- | --- | --- | --- | --- |
| 1 | Sample1_LME1 | *Streptomyces griseorubiginosus* | *Streptomyces* | CP032427.1 | 889 | 100 | 889/889 | 1 |
| 2 | Sample1_LME2 | *Streptomyces griseorubiginosus* | *Streptomyces* | CP032427.1 | 826 | 100 | 826/826 | 1 |
| 3 | Sample1_LME3 | *Rhizobium grahamii* | *Rhizobium* | PP111665.1 | 922 | 99.89 | 911/912 | 1 |
| 4 | Sample1_LME4 | *Actinacidiphila glaucinigra* (*1) | *Actinacidiphila* | CP108152.1 | 896 | 100 | 896/896 | 1 |
| 5 | Sample1_LME5 | *Streptomyces recifensis* (*2) | *Streptomyces* | MH482905.1 | 795 | 100 | 795/795 | 1 |
| 6 | Sample1_LME6 | *Actinacidiphila glaucinigra* (*1) | *Actinacidiphila* | CP108152.1 | 885 | 100 | 885/885 | 1 |
| 7 | Sample1_LME7 | *Streptomyces tanashiensis* | *Streptomyces* | CP084204.1 | 865 | 100 | 865/865 | 1 |
| 8 | Sample1_LME8 | *Streptomyces pseudovenezuelae* | *Streptomyces* | LC425853.1 | 886 | 100 | 886/886 | 1 |
| 9 | Sample1_LME9 | *Streptomyces osmaniensis* | *Streptomyces* | MT197304.1 | 639 | 100 | 639/639 | 1 |
| 10 | Sample1_LME10 | *Kitasatospora camelliae* | *Kitasatospora* | PQ097302.1 | 783 | 100 | 783/783 | 1 |
| 11 | Sample1_LME11 | *Lysobacter enzymogenes* | *Lysobacter* | OR781337.1 | 853 | 100 | 853/853 | 1 |
| 12 | Sample1_LME12 | *Rhizobium grahamii* | *Rhizobium* | PP111665.1 | 885 | 99.77 | 883/885 | 1 |
| 13 | Sample1_LME13 | *Streptomyces graminofaciens* | *Streptomyces* | AP018448.1 | 815 | 99.75 | 813/815 | 1 |
| 14 | Sample1_LME14 | *Streptomyces caeruleatus* | *Streptomyces* | KY908431.1 | 895 | 100 | 895/895 | 1 |
| 15 | Sample1_LME15 | *Streptomyces pseudovenezuelae* | *Streptomyces* | MK519177.1 | 877 | 100 | 877/877 | 1 |
| 16 | Sample1_LME16 | *Phyllobacterium brassicacearum* | *Phyllobacterium* | MT065734.1 | 957 | 100 | 957/957 | 1 |
| 17 | Sample1_LME17 | *Ramlibacter paludis* | *Ramlibacter* | MW164935.1 | 820 | 100 | 820/820 | 1 |
| 18 | Sample1_LME18 | *Rhizobium grahamii* | *Rhizobium* | PP111665.1 | 906 | 99.89 | 905/906 | 1 |
| 19 | Sample1_LME19 | *Rhizobium grahamii* | *Rhizobium* | PP111665.1 | 892 | 99.89 | 891/892 | 1 |
| 20 | Sample1_LME20 | *Kitasatospora camelliae* | *Kitasatospora* | PQ097302.1 | 877 | 100 | 877/877 | 1 |
| 21 | Sample1_LME21 | *Streptomyces tanashiensis* | *Streptomyces* | CP084204.1 | 889 | 100 | 889/889 | 1 |
| 22 | Sample1_LME22 | *Streptomyces avidinii* | *Streptomyces* | MT525247.1 | 870 | 100 | 870/870 | 1 |
| 23 | Sample1_LME23 | *Streptomyces canus* | *Streptomyces* | MT525244.1 | 887 | 99.89 | 886/887 | 1 |
| 24 | Sample1_LME24 | *Rhizobium* sp. | *Rhizobium* | OM909440.1 | 824 | 100 | 824/824 | 1 |
| 25 | Sample1_LME25 | *Actinacidiphila glaucinigra* (*1) | *Actinacidiphila* | CP108152.1 | 890 | 100 | 890/890 | 1 |
| 26 | Sample1_LME26 | *Streptomyces scabiei* | *Streptomyces* | MN372369.1 | 739 | 99.86 | 738/739 | 1 |
| 27 | Sample1_LME27 | *Rhizobium viscosum* | *Rhizobium* | MT534119.1 | 903 | 100 | 903/903 | 1 |
| 28 | Sample1_LME28 | *Streptomyces avidinii* | *Streptomyces* | MT525247.1 | 887 | 100 | 887/887 | 1 |
| 29 | Sample1_LME29 | *Rhizobium grahamii* | *Rhizobium* | PP111665.1 | 897 | 99.89 | 896/897 | 1 |
| 30 | Sample1_LME30 | *Streptomyces* sp. | *Streptomyces* | MF197388.1 | 881 | 100 | 881/881 | 1 |
| 31 | Sample1_LME31 | *Streptomyces virginiae* | *Streptomyces* | MW037167.1 | 883 | 100 | 883/883 | 1 |
| 32 | Sample1_LME32 | *Streptomyces* sp. | *Streptomyces* | CP108095.1 | 907 | 99.89 | 906/907 | 1 |
| 33 | Sample1_LME34 | *Rhizobium cauense* | *Rhizobium* | KY292451.1 | 910 | 100 | 910/910 | 1 |
| 34 | Sample1_LME35 | *Priestia megaterium* (*3) | *Priestia* | CP150971.1 | 886 | 100 | 886/886 | 1 |
| 35 | Sample1_LME36 | *Rhizobium grahamii* | *Rhizobium* | PP111665.1 | 910 | 99.89 | 909/910 | 1 |
| 36 | Sample1_LME37 | *Streptomyces chartreusis* | *Streptomyces* | CP023689.1 | 897 | 100 | 897/897 | 1 |
| 37 | Sample1_LME38 | *Streptomyces canus* | *Streptomyces* | MT525244.1 | 887 | 99.89 | 886/887 | 1 |
| 38 | Sample1_LME39 | *Streptomyces phaeofaciens* | *Streptomyces* | KU324450.1 | 873 | 100 | 873/873 | 1 |
| 39 | Sample1_LME40 | *Streptomyces* sp. | *Streptomyces* | PP837890.1 | 886 | 100 | 886/886 | 1 |
| 40 | Sample1_LME41 | *Variovorax gossypii* | *Variovorax* | OQ422116.1 | 864 | 99.88 | 863/864 | 1 |
| 41 | Sample1_LME42 | *Microvirga ossetica* | *Microvirga* | PQ870330.1 | 923 | 99.89 | 922/923 | 1 |
| 42 | Sample1_LME43 | *Pseudomonas koreensis* | *Pseudomonas* | MN272363.1 | 938 | 100 | 938/938 | 1 |
| 43 | Sample1_LME44 | *Lysobacter enzymogenes* | *Lysobacter* | OQ383329.1 | 914 | 100 | 914/914 | 1 |
| 44 | Sample1_LME45 | *Rhizobium viscosum* | *Rhizobium* | MK135797.1 | 920 | 100 | 920/920 | 1 |
| 45 | Sample1_LME46 | *Phyllobacterium pellucidum* | *Phyllobacterium* | MN658537.1 | 805 | 100 | 805/805 | 1 |
| 46 | Sample1_LME47 | *Streptomyces microflavus* | *Streptomyces* | PV110951.1 | 792 | 100 | 792/792 | 1 |
| 47 | Sample1_LME48 | *Streptomyces tanashiensis* | *Streptomyces* | CP084204.1 | 826 | 100 | 826/826 | 1 |
| 48 | Sample1_LME49 | *Streptomyces venezuelae* | *Streptomyces* | CP029197.1 | 826 | 100 | 826/826 | 1 |
| 49 | Sample1_LME50 | *Streptomyces populi* | *Streptomyces* | OP364628.1 | 824 | 99.15 | 817/824 | 1 |
| 50 | Sample1_LME51 | *Streptomyces populi* | *Streptomyces* | OP364628.1 | 824 | 99.27 | 818/824 | 1 |
| 51 | Sample1_LME52 | *Streptomyces lannensis* | *Streptomyces* | MK026800.1 | 924 | 99.46 | 919/924 | 1 |
| 52 | Sample1_LME53 | *Variovorax gossypii* | *Variovorax* | OQ422116.1 | 894 | 99.89 | 893/894 | 1 |
| 53 | Sample1_LME55 | *Ramlibacter paludis* | *Ramlibacter* | MW164935.1 | 932 | 100 | 932/932 | 1 |
| 54 | Sample1_LME56 | *Lysobacter enzymogenes* | *Lysobacter* | OQ383329.1 | 877 | 99.43 | 872/877 | 1 |
| 55 | Sample1_LME58 | *Streptomyces chartreusis* | *Streptomyces* | CP023689.1 | 838 | 100 | 838/838 | 1 |
| 56 | Sample1_LME59 | *Lysobacter prati* | *Lysobacter* | OR431958.1 | 922 | 99.89 | 921/922 | 1 |
| 57 | Sample1_LME60 | *Streptomyces canus* | *Streptomyces* | MT525246.1 | 882 | 99.21 | 875/882 | 1 |
| 58 | Sample1_LME63 | *Peristeroidobacter agariperforans* | *Peristeroidobacter* | MW186159.1 | 908 | 99.56 | 904/908 | 1 |
| 59 | Sample1_LME64 | *Streptomyces osmaniensis* | *Streptomyces* | MT197304.1 | 770 | 98.83 | 761/770 | 1 |
| 60 | Sample1_LME65 | *Streptomyces chartreusis* | *Streptomyces* | MG385711.1 | 936 | 100 | 936/936 | 1 |
| 61 | Sample1_LME66 | *Streptomyces canus* | *Streptomyces* | MT525244.1 | 916 | 99.89 | 915/916 | 1 |
| 62 | Sample1_LME67 | *Lysobacter enzymogenes* | *Lysobacter* | OQ383329.1 | 942 | 99.47 | 937/942 | 1 |
| 63 | Sample1_LME68 | *Brevundimonas alba* | *Brevundimonas* | OQ255329.1 | 871 | 99.66 | 868/871 | 1 |
| 64 | Sample1_LME70 | *Streptomyces chartreusis* | *Streptomyces* | CP023689.1 | 888 | 100 | 888/888 | 1 |
| 65 | Sample1_LME71 | *Streptomyces goshikiensis* | *Streptomyces* | MT197301.1 | 898 | 100 | 898/898 | 1 |
| 66 | Sample2_LME1 | *Streptomyces luteogriseus* | *Streptomyces* | OM980673.1 | 802 | 99.88 | 801/802 | 2 |
| 67 | Sample2_LME2 | *Streptomyces virginiae* | *Streptomyces* | MT525258.1 | 888 | 100 | 888/888 | 2 |
| 68 | Sample2_LME3 | *Actinophytocola corallina* | *Actinophytocola* | NR_112955.1<br>(*5) | 491 | 98.98 | 486/491 | 2 |
| 69 | Sample2_LME4 | *Bacillus* sp. (*4) | *Bacillus* | MT102725.1 | 894 | 100 | 894/894 | 2 |
| 70 | Sample2_LME5 | *Paenibacillus terricola* | *Paenibacillus* | MN496382.1 | 778 | 99.61 | 775/778 | 2 |
| 71 | Sample2_LME6 | *Streptomyces scabiei* | *Streptomyces* | PQ810078.1 | 899 | 99.89 | 898/899 | 2 |
| 72 | Sample2_LME7 | *Priestia megaterium* (*3) | *Priestia* | CP150971.1 | 892 | 100 | 892/892 | 2 |
| 73 | Sample2_LME8 | *Priestia aryabhattai* (*3) | *Priestia* | MZ067916.1 | 891 | 100 | 891/891 | 2 |
| 74 | Sample2_LME9 | *Streptomyces misionensis* | *Streptomyces* | MK968130.1 | 889 | 100 | 889/889 | 2 |
| 75 | Sample2_LME10 | *Priestia aryabhattai* (*3) | *Priestia* | PV478422.1 | 887 | 100 | 887/887 | 2 |
| 76 | Sample2_LME11 | *Streptomyces akebiae* | *Streptomyces* | CP080647.1 | 890 | 100 | 890/890 | 2 |
| 77 | Sample2_LME13 | *Streptomyces luteogriseus* | *Streptomyces* | OM980673.1 | 890 | 100 | 890/890 | 2 |
| 78 | Sample2_LME14 | *Paenibacillus xanthanilyticus* | *Paenibacillus* | NR_159286.1<br>(*5) | 821 | 99.63 | 828/821 | 2 |
| 79 | Sample2_LME15 | *Streptomyces lincolnensis* | *Streptomyces* | KU870779.1 | 890 | 99.55 | 886/890 | 2 |
| 80 | Sample2_LME16 | *Streptomyces neyagawaensis* | *Streptomyces* | NR_025868.2<br>(*5) | 890 | 100 | 890/890 | 2 |
| 81 | Sample2_LME17 | *Streptomyces neyagawaensis* | *Streptomyces* | NR_025868.2<br>(*5) | 827 | 100 | 827/827 | 2 |
| 82 | Sample2_LME18 | *Priestia aryabhattai* (*3) | *Priestia* | PV478422.1 | 887 | 100 | 887/887 | 2 |
| 83 | Sample2_LME19 | *Actinophytocola corallina* | *Actinophytocola* | NR_112955.1<br>(*5) | 888 | 98.87 | 878/888 | 2 |
| 84 | Sample2_LME20 | *Streptomyces misionensis* | *Streptomyces* | MK968130.1 | 889 | 100 | 889/889 | 2 |
| 85 | Sample2_LME21 | *Bacillus* sp. (*4) | *Bacillus* | MT102725.1 | 894 | 100 | 894/894 | 2 |
| 86 | Sample2_LME22 | *Bacillus* sp. (*4) | *Bacillus* | MT102725.1 | 894 | 100 | 894/894 | 2 |
| 87 | Sample2_LME23 | *Actinophytocola corallina* | *Actinophytocola* | NR_112955.1<br>(*5) | 591 | 99.15 | 586/591 | 2 |
| 88 | Sample2_LME25 | *Actinophytocola corallina* | *Actinophytocola* | NR_112955.1<br>(*5) | 497 | 98.99 | 492/497 | 2 |
| 89 | Sample2_LME26 | *Bradyrhizobium lablabi* | *Bradyrhizobium* | LT670844.1 | 898 | 100 | 898/898 | 2 |
| 90 | Sample2_LME27 | *Streptomyces luteogriseus* | *Streptomyces* | OM980673.1 | 519 | 100 | 519/519 | 2 |
| 91 | Sample2_LME28 | *Actinophytocola corallina* | *Actinophytocola* | NR_112955.1<br>(*5) | 648 | 99.23 | 643/648 | 2 |
| 92 | Sample3_LME1 | *Pinibacter aurantiacus* | *Pinbacter* | MH368770.2 | 873 | 99.54 | 869/873 | 3 |
| 93 | Sample3_LME2 | *Streptomyces bungoensis* | *Streptomyces* | MT124448.1 | 367 | 100 | 367/367 | 3 |
| 94 | Sample3_LME3 | *Rhizobium* sp. | *Rhizobium* | KY445677.1 | 887 | 100 | 887/887 | 3 |
| 95 | Sample3_LME4 | *Streptomyces kronopolitis* | *Streptomyces* | ON819565.1 | 910 | 100 | 910/910 | 3 |
| 96 | Sample3_LME5 | *Uncultured Dyella* sp. | *Dyella* | PP780941.1 | 907 | 100 | 907/907 | 3 |
| 97 | Sample3_LME6 | *Streptomyces mirabilis* | *Streptomyces* | MK254602.1 | 824 | 100 | 824/824 | 3 |
| 98 | Sample3_LME7 | *Paraburkholderia caribensis* | *Paraburkholderia* | MG270180.1 | 845 | 100 | 845/845 | 3 |
| 99 | Sample3_LME8 | *Kitasatospora herbaricolor* (*1) | *Kitasatospora* | MT081107.1 | 889 | 100 | 889/889 | 3 |
| 100 | Sample3_LME9 | *Streptomyces kronopolitis* | *Streptomyces* | ON819565.1 | 908 | 100 | 908/908 | 3 |
| 101 | Sample3_LME10 | *Kitasatospora herbaricolor* (*1) | *Kitasatospora* | MT081107.1 | 150 | 100 | 150/150 | 3 |
| 102 | Sample3_LME11 | *Buttiauxella* sp. | *Buttiauxella* | CP110080.1 | 850 | 98.94 | 841/850 | 3 |
| 103 | Sample3_LME12 | *Caulobacter* sp. | *Caulobacter* | JQ977670.1 | 884 | 100 | 884/884 | 3 |
| 104 | Sample3_LME13 | *Neorhizobium xiangyangii* | *Neorhizobium* | MT611284.1 | 765 | 99.22 | 759/765 | 3 |
| 105 | Sample3_LME14 | *Uncultured Caulobacter* sp. | *Caulobacter* | JQ977670.1 | 914 | 100 | 914/914 | 3 |
| 106 | Sample3_LME15 | *Streptomyces* sp. | *Streptomyces* | CP108774.1 | 889 | 99.89 | 888/889 | 3 |
| 107 | Sample3_LME16 | *Streptomyces* sp. | *Streptomyces* | OQ155109.1 | 387 | 99.74 | 386/387 | 3 |
| 108 | Sample3_LME17 | *Streptomyces* sp. | *Streptomyces* | KX023623.1 | 300 | 100 | 300/300 | 3 |
| 109 | Sample3_LME18 | *Arthrobacter globiformis* | *Arthrobacter* | ON838146.1 | 872 | 99.77 | 870/872 | 3 |
| 110 | Sample3_LME19 | *Uncultured Dyella* sp. | *Dyella* | PP780941.1 | 906 | 100 | 906/906 | 3 |
| 111 | Sample3_LME20 | *Labrys miyagiensis* | *Labrys* | AB681465.1 | 917 | 100 | 917/917 | 3 |
| 112 | Sample3_LME21 | *Phenylobacterium* sp. | *Phenylobacterium* | OR878925.1 | 905 | 99.89 | 904/905 | 3 |
| 113 | Sample3_LME22 | *Nocardoides* sp. | *Nocardoides* | LN876367.1 | 125 | 100 | 125/125 | 3 |
| 114 | Sample3_LME24 | *Weissella paramesenteroides* | *Weissella* | PV270261.1 | 698 | 99.57 | 695/698 | 3 |
| 115 | Sample3_LME25 | *Conexibacter* sp. | *Conexibacter* | MK875928.1 | 907 | 99.01 | 898/907 | 3 |
| 116 | Sample3_LME26 | *Mesorhizobium qingshengii* | *Mesorhizobium* | PV018831.1 | 895 | 100 | 895/895 | 3 |
| 117 | Sample3_LME27 | *Labrys* sp. | *Labrys* | OQ647604.1 | 898 | 100 | 898/898 | 3 |
| 118 | Sample3_LME28 | *Cellulomonas* sp. | *Cellulomonas* | OQ346295.1 | 660 | 99.85 | 659/660 | 3 |
| 119 | Sample3_LME29 | *Rhizobium* sp. | *Rhizobium* | LC372620.1 | 656 | 100 | 656/656 | 3 |
| 120 | Sample3_LME30 | *Caballeronia* sp. | *Caballeronia* | OR373818.1 | 746 | 99.46 | 742/746 | 3 |
| 121 | Sample3_LME31 | *Nocardoides* sp. | *Nocardoides* | JQ977357.1 | 860 | 99.77 | 858/860 | 3 |
| 122 | Sample3_LME34 | *Cupriavidus* sp. | *Cupriavidus* | KY381884.1 | 908 | 99.01 | 899/908 | 3 |
| 123 | Sample3_LME35 | *Silvimonas terrae* | *Silvimonas* | NR_113961.1 | 144 | 100 | 144/144 | 3 |
| 124 | Sample3_LME36 | *Mesorhizobium qingshengii* | *Mesorhizobium* | PV018831.1 | 895 | 100 | 895/895 | 3 |
| 125 | Sample3_LME37 | *Dyella japonica* | *Dyella* | OP102632.1 | 876 | 99.89 | 875/876 | 3 |
| 126 | Sample3_LME38 | *Weissella paramesenteroides* | *Weissella* | JN863617.1 | 644 | 99.84 | 644/645 | 3 |
| 127 | Sample3_LME39 | *Labrys monachus* | *Labrys* | MT534155.1 | 600 | 99.5 | 597/600 | 3 |

| No. | Strain Name | Most closely related species | Genus | GenBank<br>accession No. | Read<br>length<br>(bp) | Identity<br>(%) | Identical<br>base pair | Sample |
| --- | --- | --- | --- | --- | --- | --- | --- | --- |
| 128 | Sample1_LME33 | *Duganella* sp. | *Not determined* | JQ745646.1 | 837 | 98.69 | 826/837 | 1 |
| 129 | Sample1_LME54 | *Paraflavitalea speifideaquila* | *Not determined* | CP135920.1 | 367 | 98.09 | 360/367 | 1 |
| 130 | Sample1_LME57 | *Uncultured Ohtaekwangia* sp. | *Not determined* | JQ771968.1 | 954 | 97.79 | 931/952 | 1 |
| 131 | Sample1_LME61 | *Ramlibacter monticola* | *Not determined* | PP595918.1 | 867 | 98,27 | 852/867 | 1 |
| 132 | Sample1_LME62 | *Niastella* sp. | *Not determined* | MH490984.1 | 890 | 96.77 | 839/867 | 1 |
| 133 | Sample1_LME69 | *Niastella* sp. | *Not determined* | MH490984.1 | 931 | 96,93 | 883/911 | 1 |
| 134 | Sample2_LME12 | *Uncultured Cohnella* sp. | *Not determined* | KJ749163.1 | 279 | 97.49 | 272/279 | 2 |
| 135 | Sample2_LME24 | *Actinophytocola corallina* | *Not determined* | NR_112955.1<br>(*5) | 219 | 97.72 | 214/219 | 2 |
| 136 | Sample3_LME23 | *Labrys* sp. | *Not determined* | OQ647604.1 | 245 | 98.37 | 241/245 | 3 |
| 137 | Sample3_LME32 | *Catenulispora* sp. strain | *Not determined* | PQ637139.1 | 149 | 97.32 | 145/149 | 3 |
| 138 | Sample3_LME33 | *Jatrophihabitans* sp. | *Not determined* | MK875924.1 | 262 | 95.38 | 248/260 | 3 |

## Reference

1. Waksman Selman A, Reilly HC, Johnstone Donald B. 1946. Isolation of Streptomycin-producing Strains of *Streptomyces griseus*. Journal of Bacteriology 52:393–397.

2. Waksman SA. 1953. Streptomycin: background, isolation, properties, and utilization. Science 118:259–66.

3. Tomishima M, Ohki H, Yamada A, Takasugi H, Maki K, Tawara S, Tanaka H. 1999. FK463, a novel water-soluble echinocandin lipopeptide: synthesis and antifungal activity. J Antibiot (Tokyo) 52:674–6.

4. Burg RW, Miller BM, Baker EE, Birnbaum J, Currie SA, Hartman R, Kong YL, Monaghan RL, Olson G, Putter I, Tunac JB, Wallick H, Stapley EO, Oiwa R, Omura S. 1979. Avermectins, new family of potent anthelmintic agents: producing organism and fermentation. Antimicrob Agents Chemother 15:361–7.

5. Kino T, Hatanaka H, Hashimoto M, Nishiyama M, Goto T, Okuhara M, Kohsaka M, Aoki H, Imanaka H. 1987. FK-506, a novel immunosuppressant isolated from a *Streptomyces*. I. Fermentation, isolation, and physico-chemical and biological characteristics. J Antibiot (Tokyo) 40:1249–55.

6. Arcamone F, Cassinelli G, Fantini G, Grein A, Orezzi P, Pol C, Spalla C. 1969. Adriamycin, 14-hydroxydaunomycin, a new antitumor antibiotic from *S. peucetius* var. caesius. Biotechnol Bioeng 11:1101–10.

7. Hamamoto H, Urai M, Ishii K, Yasukawa J, Paudel A, Murai M, Kaji T, Kuranaga T, Hamase K, Katsu T, Su J, Adachi T, Uchida R, Tomoda H, Yamada M, Souma M, Kurihara H, Inoue M, Sekimizu K. 2015. Lysocin E is a new antibiotic that targets menaquinone in the bacterial membrane. Nat Chem Biol 11:127–33.

8. Hamamoto H, Panthee S, Paudel A, Ishii K, Yasukawa J, Su J, Miyashita A, Itoh H, Tokumoto K, Inoue M, Sekimizu K. 2021. Serum apolipoprotein A-I potentiates the therapeutic efficacy of lysocin E against *Staphylococcus aureus*. Nat Commun 12:6364.

9. van der Heijden MG, Bardgett RD, van Straalen NM. 2008. The unseen majority: soil microbes as drivers of plant diversity and productivity in terrestrial ecosystems. Ecol Lett 11:296–310.

10. Gans J, Wolinsky M, Dunbar J. 2005. Computational improvements reveal great bacterial diversity and high metal toxicity in soil. Science 309:1387–90.

11. Kato S, Yamagishi A, Daimon S, Kawasaki K, Tamaki H, Kitagawa W, Abe A, Tanaka M, Sone T, Asano K, Kamagata Y. 2018. Isolation of Previously Uncultured Slow-Growing Bacteria by Using a Simple Modification in the Preparation of Agar Media. Applied and Environmental Microbiology 84:e00807–18.

12. Harwani D. 2013. The great plate count anomaly and the unculturable bacteria. Microbiology 2:350–1.

13. Hazu M, Ahmed A, Curry E, Hornby DP, Gjerde DT. 2022. Cell Chromatography: Biocompatible Chromatographic Separation and Interrogation of Microbial Cells. Microbiol Spectr 10:e0245022.

14. Taha MPM, Drew GH, Tamer Vestlund A, Aldred D, Longhurst PJ, Pollard SJT. 2007. Enumerating actinomycetes in compost bioaerosols at source—Use of soil compost agar to address plate ‘masking’. Atmospheric Environment 41:4759–4765.

15. Janssen PH, Yates PS, Grinton BE, Taylor PM, Sait M. 2002. Improved culturability of soil bacteria and isolation in pure culture of novel members of the divisions Acidobacteria, Actinobacteria, Proteobacteria, and Verrucomicrobia. Appl Environ Microbiol 68:2391–6.

16. Bloomfield SF, Stewart G, Dodd CER, Booth IR, Power EGM. 1998. The viable but non-culturable phenomenon explained? Microbiology (Reading) 144 (Pt 1):1–3.

17. Aldén L, Demoling F, Bååth E. 2001. Rapid Method of Determining Factors Limiting Bacterial Growth in Soil. Applied and Environmental Microbiology 67:1830–1838.

18. Tanaka Y, Hanada S, Manome A, Tsuchida T, Kurane R, Nakamura K, Kamagata Y. 2004. Catellibacterium nectariphilum gen. nov., sp. nov., which requires a diffusible compound from a strain related to the genus Sphingomonas for vigorous growth. International Journal of Systematic and Evolutionary Microbiology 54:955–959.

19. Goh CBS, Goh CHP, Wong LW, Cheng WT, Yule CM, Ong KS, Lee SM, Pasbakhsh P, Tan JBL. 2022. A three-dimensional (3D) printing approach to fabricate an isolation chip for high throughput in situ cultivation of environmental microbes. Lab Chip 22:387–402.

20. Ho YN, Chen YL, Liu DY. 2021. Portable and Rapid Sequencing Device with Microbial Community-Guided Culture Strategies for Precious Field and Environmental Samples. mSystems 6:e0074821.

21. Stapley EO, Jackson M, Hernandez S, Zimmerman SB, Currie SA, Mochales S, Mata JM, Woodruff HB, Hendlin D. 1972. Cephamycins, a new family of beta-lactam antibiotics. I. Production by actinomycetes, including Streptomyces lactamdurans sp. n. Antimicrob Agents Chemother 2:122–31.

22. Miyashita A, Kataoka K, Tsuchida T, Ogasawara AA, Nakajima H, Takahashi M, Sekimizu K. 2023. High molecular weight glucose homopolymer of broccoli (*Brassica oleracea* var. *italica*) stimulates both invertebrate and mammalian immune systems. Frontiers in Food Science and Technology 3.

23. Miyashita A, Sekimizu K, Kaito C. 2022. Surrounding gas composition affects the calling song development in the two-spotted cricket (*Gryllus bimaculatus*). Drug Discoveries & Therapeutics advpub.

24. Miyashita A, Mitsutomi S, Mizushima T, Sekimizu K. 2022. Repurposing the PDMA-approved drugs in Japan using an insect model of staphylococcal infection. FEMS Microbes 3:xtac014.

25. Miyashita A, Sekimizu K. 2021. Using silkworms to search for lactic acid bacteria that contribute to infection prevention and improvement of hyperglycemia. Drug Discoveries & Therapeutics 15:51–54.

26. Miyashita A, Hamamoto H, Sekimizu K. 2021. Applying the silkworm model for the search of immunosuppressants. Drug Discoveries & Therapeutics 15:139–142.

27. Miyashita A, Kizaki H, Sekimizu K, Kaito C. 2016. Body-enlarging effect of royal jelly in a non-holometabolous insect species, *Gryllus bimaculatus*. Biology Open 5:770–776.

28. Miyashita A, Kizaki H, Sekimizu K, Kaito C. 2016. No Effect of Body Size on the Frequency of Calling and Courtship Song in the Two-Spotted Cricket, *Gryllus bimaculatus*. PLoS One 11:e0146999.

29. Miyashita A, Takahashi S, Ishii K, Sekimizu K, Kaito C. 2015. Primed Immune Responses Triggered by Ingested Bacteria Lead to Systemic Infection Tolerance in Silkworms. PLOS ONE 10:e0130486.

30. Miyashita A, Hirai Y, Sekimizu K, Kaito C. 2015. Antibiotic-producing bacteria from stag beetle mycangia. Drug Discoveries & Therapeutics advpub.

31. Miyashita A, Kizaki H, Kawasaki K, Sekimizu K, Kaito C. 2014. Primed Immune Responses to Gram-negative Peptidoglycans Confer Infection Resistance in Silkworms *. Journal of Biological Chemistry 289:14412–14421.

32. Miyashita A, Iyoda S, Ishii K, Hamamoto H, Sekimizu K, Kaito C. 2012. Lipopolysaccharide O-antigen of enterohemorrhagic *Escherichia coli* O157:H7 is required for killing both insects and mammals. FEMS Microbiology Letters 333:59–68.

33. Kaeberlein T, Lewis K, Epstein SS. 2002. Isolating “uncultivable” microorganisms in pure culture in a simulated natural environment. Science 296:1127–9.

34. Aoi Y, Kinoshita T, Hata T, Ohta H, Obokata H, Tsuneda S. 2009. Hollow-Fiber Membrane Chamber as a Device for In Situ Environmental Cultivation. Applied and Environmental Microbiology 75:3826–3833.

35. Hayakawa M, Nonomura H. 1987. Humic acid-vitamin agar, a new medium for the selective isolation of soil actinomycetes. Journal of Fermentation Technology 65:501–509.

36. Stackebrandt E, and Ebers, J. 2006. Taxonomic parameters revisited: tarnished gold standards. Microbiology Today 33:152–155.

